# Reconciling Cooperativity Definition in PROTACs and Molecular Glues: Thermodynamic Dissection into PPI and Ligand Entropic Contributions

**DOI:** 10.64898/2026.02.07.704601

**Authors:** Yang Li, Zuohang Qu, Lizhi Jiang, Lijing Tang, Haomin Chen, Xuan Cao

**Affiliations:** Institute of Pharmacy and Pharmacology, Hunan Province, Cooperative Innovation Center for Molecular Target New Drug Study, College of Pharmacy, Hengyang Medical School, University of South China, Hengyang, Hunan, 421001, China; School of Computer Science and Engineering, Central South University, Changsha, China; Hunan Provincial Key Laboratory of Basic and Clinical Pharmacological Research of Gastrointestinal Cancer, Department of Pharmacy, Institute of Pharmacy and Pharmacology, The Second Affiliated Hospital, Hengyang Medical School, University of South China, Hengyang, Hunan, 421001, China; Hubei Bio-Pharmaceutical Industrial Technological Institute Inc., Humanwell Healthcare Co. Ltd., Wuhan, Hubei, China

**Author notes:** Corresponding Author: College of Pharmacy, Hengyang Medical School, University of South China, Hengyang, Hunan 421001, PR China, E-mail addresses (Dr. X. Cao). the authors contribute equally to this work (Yang Li, Zuohang Qu, Lizhi Jiang, Lijing Tang).

**Keywords:** Cooperativity, PROTAC, Molecular Glue, Ligand Entropy

## Abstract

Douglass’ Cooperativity *α* and Ciulli’s Cooperativity *α* in induced-proximity systems, remains controversial with paradoxes such as path-dependent metrics and apparent universal negative Cooperativity. We noticed that in “partial-embedded” model, a substantial portion of giant ligand remains exposed outside and does not engage with the host protein’s force field. It incurs an entropic cost due to the restriction of translational/rotational degrees of freedom. This large, mass-dependent unfavorable ligand entropy penalty normally shifts binding affinity to 10^4^∼10^8^-fold. ITC thermodynamic cycles analysis confirmed the dramatic entropy loss among reaction pair. This reconciles the conflicting Cooperativity definitions, yielding true path-independent positive PPI Cooperativity from observed entropy loss subtracting ligand entropy penalty. ITC data showed rigid linkers appear superior to flexible linkers with respect to both oral bioavailability and safety profile in PROTAC design. “ligand entropy barrier wall**/**Cooperativity ladder” pair is not only impact induced-proximity systems but also constitute the physical basis for all biosystems.

## INTRODUCTION

Targeted protein degradation using PROTACs and molecular glues relies on the induced formation of a ternary complexes between a protein of interest (POI), a bifunctional (or monofunctional) recruiter, and an E3 ubiquitin ligase.^1-6^ The stability and degradation efficiency of these complexes are critically governed by cooperativity — the extent to which formation of the ternary complex is energetically more (or less) favorable than the sum of the corresponding binary interactions.^7-13^

In 2013, Douglass, Spiegel and co-workers introduced a rigorous mathematical framework for three-body binding equilibria and defined cooperativity α as the ratio of binding constants for ligand versus binary-complex binding to the same target.^14^ Under this classical definition, α < 1, α = 1, and α > 1 correspond to positive, zero, and negative Cooperativity, respectively. Surprisingly, virtually all reported PROTACs and molecular glues exhibit α ≫ 1 (typically 10^4^–10^8^), implying pervasive negative cooperativity or steric clash. This observation has remained puzzling, given ample structural evidence of favorable POI–E3 contacts in ternary complexes.

In 2017, Ciulli and co-workers proposed an alternative definition, as the ratio of dissociation constants for binary-complex versus ternary complex, in which α < 1 indicates enhanced ternary complex stability relative to the binary complex^15^. This empirically oriented metric rapidly became the field standard because high α values reliably predict potent cellular degradation^13,16-18^. However, the two definitions are mutually incompatible: a molecule can simultaneously appear strongly negatively cooperative (classical α ≫ 1) yet highly positively cooperative (Ciulli’s α ≪ 1). Moreover, Ciulli’s definition can give the impression that ternary complex stability is path-dependent, a notion seemingly at odds with thermodynamic cycle closure and microscopic reversibility. These inconsistencies have persisted for nearly a decade, despite widespread use of Ciulli’s α as the central design parameter in proximity-based pharmacology. ^8,13,19-22^

Herein, we resolve this long-standing paradox by showing that both definitions are simultaneously correct but measure physically distinct phenomena. We noticed that “partial-embedded” model of bifunctional molecules such as PROTAC, molecular glue or its binary complex, were quite different with traditional “full-embedded” binding model for small molecules. In “partial-embedded” model, a substantial portion of giant ligand remains exposed outside and does not engage with the host protein’s force field, it incurs an entropic cost due to the restriction of translational/rotational degrees of freedom. This large, mass-dependent unfavorable ligand entropic penalty (Δ*S*_*ligand*_) term, typically +4 to +8 kcal·mol^−1^, quantitatively accounts for the ubiquitous α ≫ 1 observed under the classical definition, eliminates apparent path dependence, and reconciles the empirical success of Ciulli’s α with rigorous thermodynamics. Our framework further reveals that true positive cooperativity in most degraders is predominantly entropic in origin, scales with induced POI– E3 interface area and is modulated by linker rigidity through enthalpy–entropy compensation. These insights not only settle a decade of confusion but also provide clear physicochemical guidelines for the rational optimization of next-generation proximity-inducing therapeutics.

### Two Definition of Cooperativity *α* and Paradox

Douglass’ Cooperativity *α* was defined as the ratio (*α* = K_d_^binary^/K_d_^ligand^). Surprisingly, virtually all reported PROTACs and molecular glues exhibit α ≫ 1 (typically 104–108), implying pervasive negative cooperativity or steric clash.

We retrieved all PROTAC reports from PROTAC-DB 3.0 database and obtained 64 records containing core parameters. These calculated Cooperativity *α* results were listed in Supplementary Table S1, which showed that all 64 records of Cooperativity *α* were all decreased without any exceptions, ranging from 20,000 to 100,000,000-fold. According to the definition of Cooperativity, that means strongly compulsive or steric hinderance occurs when ternary complex is formed. However, Cryo-EM show favorable POI–E3 contacts in ternary complexes rather than pervasive negative cooperativity or steric clash (Figure S1, Table S2). In 2017, Ciulli and co-workers proposed an alternative definition, (*α* = K_d_^ternary^/K_d_^binary^), in which α < 1 indicates enhanced ternary complex stability relative to the binary complex_15_. This empirically oriented metric rapidly became the field standard because high α values reliably predict potent cellular degradation^13,16-18^.

However, as a ternary dynamic equilibrium system, if Ciulli’s *α* represents the cooperation factor from binary to ternary complex, theoretically, as a mutual synergic process, this Cooperativity should be path-independent. And different binding sequence of binary complex would lead to dramatically different *K*_*d*_ value. Thus, it seems at odds with thermodynamic cycle closure and microscopic reversibility. We checked out PROTACDB 3.0, calculated all Cooperativity *α* in different path forming ternary, and found no report data supporting path-independence. The results were shown in Table S3.

We noticed that “partial-embedded” model of bifunctional molecules such as PROTAC, molecular glue or its binary complex, were quite different with traditional “full-embedded” binding model for small molecules. In “partial-embedded” model, a substantial portion of giant ligand remains exposed outside and does not engage with the host protein’s force field, it incurs an entropic cost due to the restriction of translational/rotational degrees of freedom (Figure 1). This unfavorable ligand entropy penalty might be the core paradox of Douglass’ Cooperativity *α* and Ciulli’s Cooperativity *α*. Thus, to calculate the Cooperativity, we should calculate these ligand entropy penalty first.

**Figure 1.**
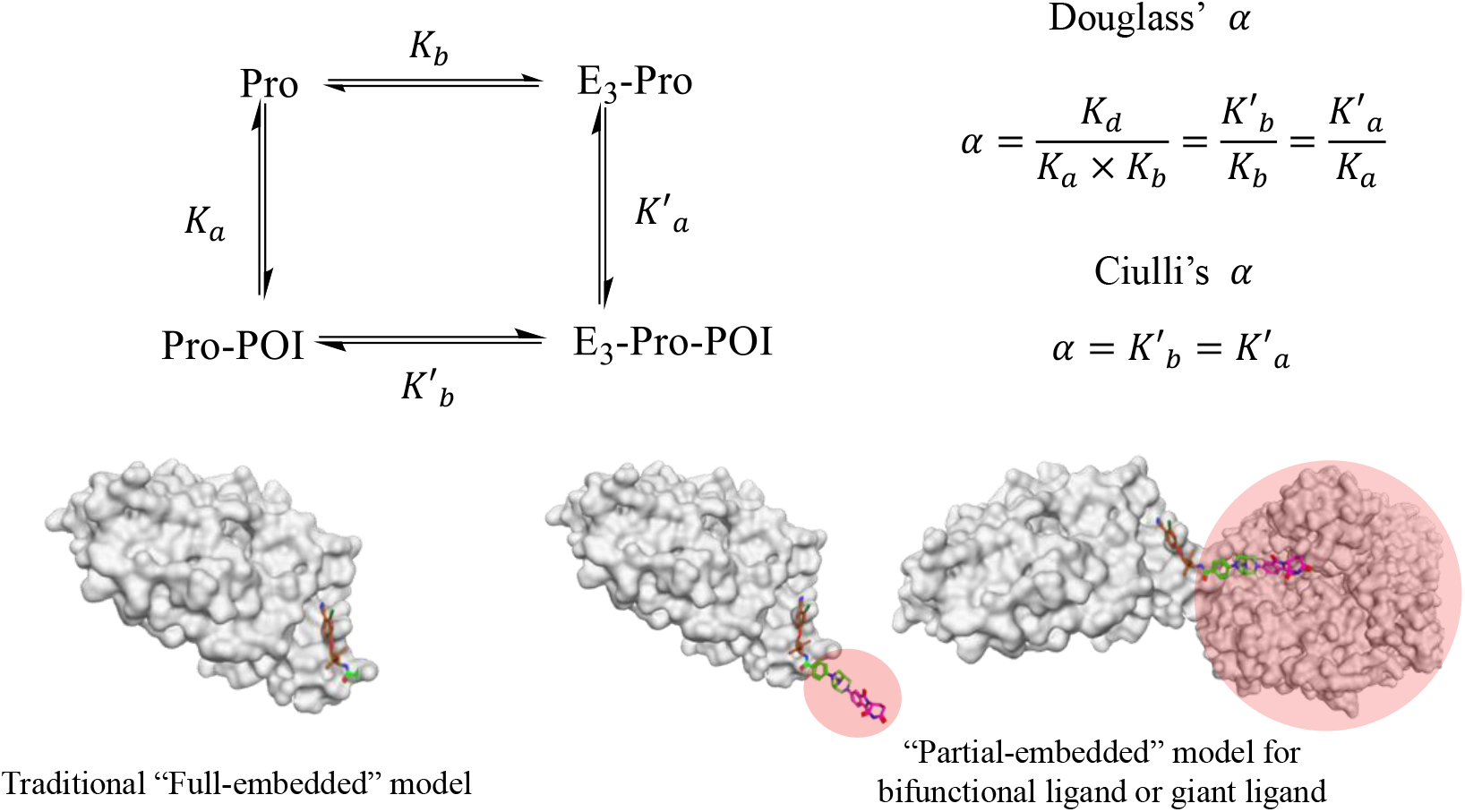
The upper shows scheme of different Cooperative definition by Douglass and Ciulli. The bottom left shows the difference between tradition “full embedded” binding model for small molecules and bottom right shows the “partial embedded” binding model for bifunctional molecules such as PROTAC and molecular glue. The pink area means the portion of giant ligand remains exposed outside and does not engage with the host protein’s force field; it incurs an entropic cost due to the restriction of translational/rotational degrees of freedom. Apparently, the giant ligand exposed part should pay more entropy penalty than small one.

### Thermodynamic Calculation of Ligand Entropy (Translational Entropy)

The translational entropy of ideal gas was first given by Sackur-Tetrode equation in 1912:^23^

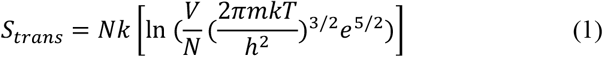

Here m refers to the mass of the ligand. Upon binding, this leads to a mass-dependent entropy loss, as formalized by Finkelstein and Janin (1989).^24^ They derived the unfavorable translational entropy change upon association (at standard conditions, 298 K) as approximately +0.9 kcal/mol for three degrees of freedom (∼+0.3 kcal/mol per degree), plus contributions from rotation as function (2).

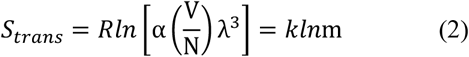

Here α=67.84, and 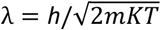. This mass dependence arises fundamentally from equipartition in classical statistical mechanics (Newton’s laws applied thermally): when average kinetic energy is fixed (1/2•mV^2^= constant), velocity 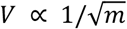 and momentum 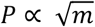. A 4-fold mass would double momentum, providing an intuitive scaling: *P* = log_2_(m’/m) = 2 (per 4-fold mass). That means heavier “giant” ligands show higher-momentum motion in free state and thermodynamic means a bigger entropic hit when constrained.

For “partial-embedded” cases, the penalty scales with the effective mass ratio M’/M (∼3 translational and rotational degree, scaled from Finkelstein–Janin):

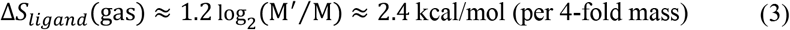

Accounting for solvent (water) effects, we empirically adjust a coefficient 0.85 to formula 3, then we get the Δ*S*_*ligand*_ function in water by function 4:

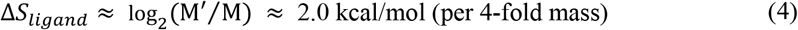

For PROTAC, M refers to the embedded molecular weight of ligand, and M’ represents the molecular weight of PROTAC. For PROTAC-POI binary complex, M refers to the molecular weight of PROTAC, and M’ represents the molecular weight of binary complex. We note that the calculation method only suitable for these “partial embedded” model which was quite different with traditional “full embedded” model.

### Disserting Ciulli’s Cooperativity *α* and Douglass’ *α*

We then elucidate Douglass’ *α* by thermodynamic theory and propose it was comprised of two distinct components. Note that when warhead binding affinity is included, Douglass’ *α* would be equal to Ciulli’s Cooperativity *α*. The first component log[*α*]_*PPI*_, arises from all energetic contributions associated with formation of the ternary complex and can be further decomposed into two terms: conformational entropy penalty (Δ*S*_*conf*_), and enthalpic contribution (ΔΔ*H*) which equals entropy–enthalpy compensation term (Δ*S*_*comp*_). The second component Δ*S*_*ligand*_, represents an additional ligand entropy penalty that is typically not accounted for in standard thermodynamic model.

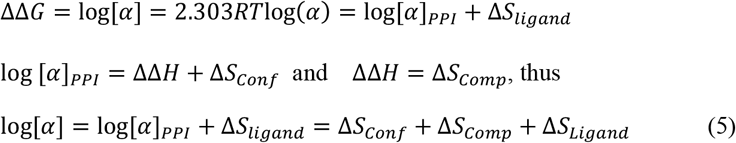

Or we can transferred the energy related term log[*α*]_*PPI*_ and Δ*S*_*ligand*_ to dissociation constant related parameter log (*α*)_*Ligand*_ and log (*α*)_*PPI*_

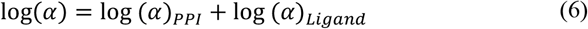

Then we could get results that when log[*α*]_*PPI*_ or log(*α*)_*PPI*_ > 0, = 0, < 0 represent negative energy binding, non-energy binding, positive energy binding, respectively.

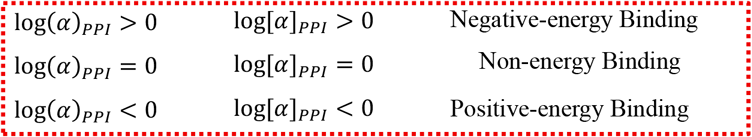

The calculation for each thermodynamic terms for contributions of reaction pair were listed in Table 1. Some cases of calculated log (*α*)_*PPI*_ were listed in Table 2. We found log(*α*)_*PPI*_ were consisted of positive, negative and non-energy releasing binding in calculation of random samples. Our theory confirmed the releasing of molecular potential energy during ternary complex forming which was dominated by ligand-entropy. That seems to correlate the paradox of Douglass’ *α* and Ciulli’s *α* after subtracting the ligand entropy penalty. However, we noted that log[*α*]_*PPI*_ still can’t represent the Cooperativity behavior of protein-protein interface, just a whole energy summary or binding affinity alteration.

**Table 1.** The calculation method of thermodynamic terms for contribution of reaction pair.

| Thermodynamics | Calculation function | Represents contribution |
| --- | --- | --- |
| $\Delta\Delta G_{\text{observed}}$ | $= (\Delta H' - T\Delta S') - (\Delta H - T\Delta S)$ | Total energy loss of the reaction pair (kcal/mol) |
| $\Delta\Delta H_{\text{observed}}$ | $= \Delta H' - \Delta H$ | Total enthalpy loss of the reaction pair (kcal/mol) |
| $\Delta\Delta S_{\text{observed}}$ | $= -T\Delta S' - (-T\Delta S)$ | Total entropy loss of the reaction pair (kcal/mol) |
| $\log(\alpha)_{\text{Ligand}}$ | $= 0.85\log_2(M'/M)$ | Ligand entropy penalty related dissociate constant $K_d$ |
| $\Delta S_{\text{ligand}}$ | $= 2.303RT \cdot 0.85\log_2(M'/M)$<br>$\approx +2$ (kcal/mol) per 4-fold Mass | Ligand entropy penalty of the reaction pair (kcal/mol) |
| $\log(\alpha)_{\text{PPI}}$ | $= \log(K_d) - \log(K_a \cdot K_b) - 0.85\log_2(M'/M)$ | Total related dissociate constant $K_d$ change in formation of ternary complex |
| $\log[\alpha]_{\text{PPI}}$ | $= \Delta\Delta G_{\text{observed}} - \Delta S_{\text{ligand}}$ | Total energy change in formation of ternary complex (kcal/mol) |
| $\Delta S_{\text{Comp}}$ | $= \Delta\Delta H_{\text{observed}}$ | Compensation for the conformation change of PROTAC (kcal/mol) |
| $\Delta S_{\text{Conf}}$ | $= \Delta\Delta S_{\text{observed}} - \Delta S_{\text{ligand}}$ | Real Cooperativity for PPI interface (kcal/mol) |

**Table 2.**
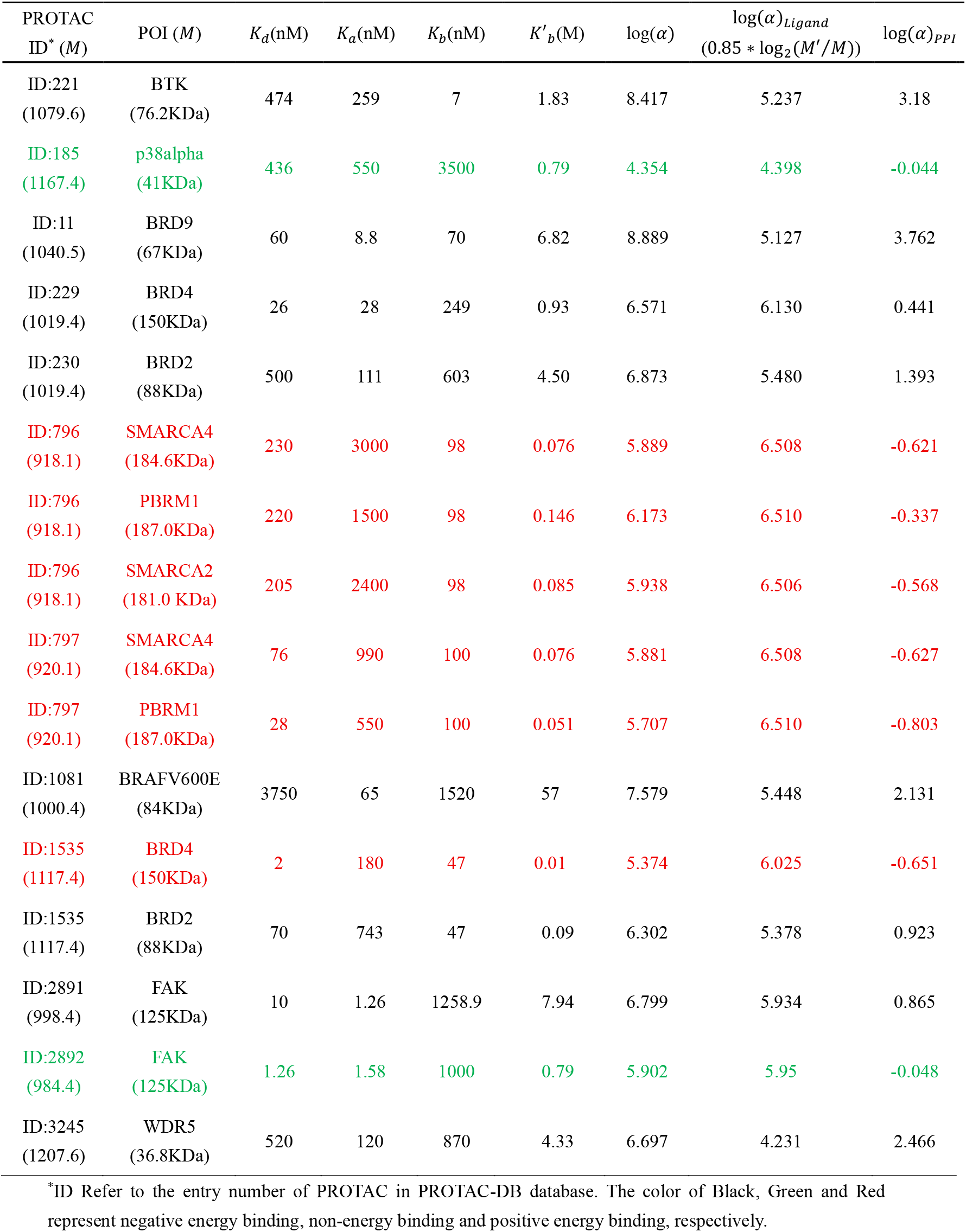
Calculation of Cooperativity log(*α*), log(*α*)_*PPI*_, log (*α*)_*Ligand*_ of PROTAC.

### Isothermal Titration Calorimetry (ITC) Data Analysis

Isothermal titration calorimetry (ITC) was used to dissect the complete thermodynamic cycle governing ternary complex formation. In 2025, *Amit Vaish* et al. utilized ITC to interrogate the hydrodynamic and thermodynamic parameters influencing molecular dynamics and ternary complex stability in solution.^25^ Specifically, they examined the sequential assembly of binary and ternary complexes involving the PROTAC hetero-bifunctional small molecule (hSM; compound 1), VHL-ElonginC-ElonginB (VBC), and SMARCA2 (SM2) or SMARCA4 (SM4) as the protein of interest (POI). The original data were listed in Table S4. However, the full cycle data was apparently path-dependent for ternary complex and seemingly at odds with thermodynamic cycle closure and microscopic reversibility. Thus, we correct the ITC cycle data and list it in Table S5. The analysis of these corrected titration results was summarized in Figure 2.

**Figure 2.**
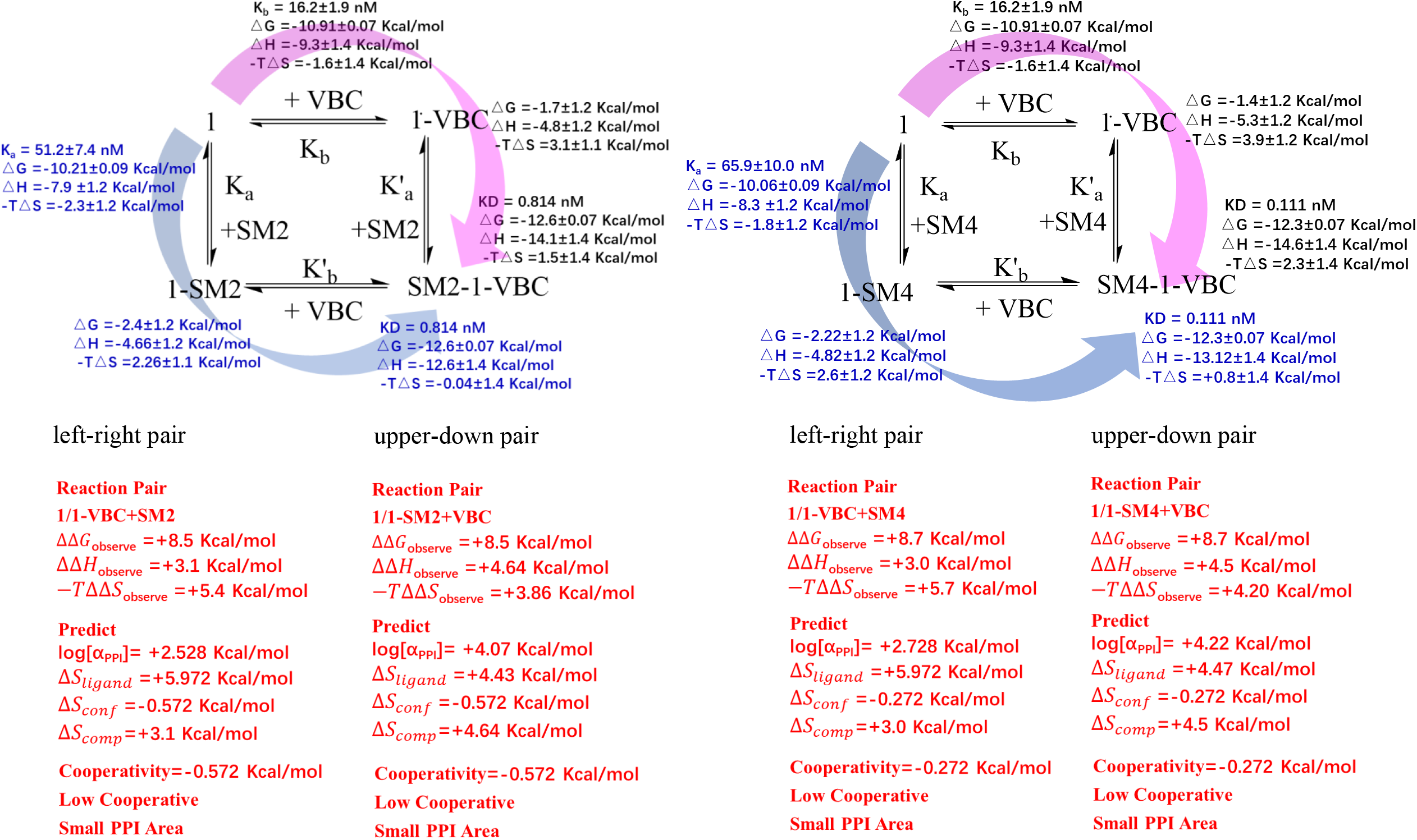
Revised full ITC thermodynamic cycle data of binary/ternary complex from Compound 1, VBC, and SM2/SM4.

Holly Soutter’s group^26^ titrates the energy contributions in reaction pair of MZ1 /MZ1-BRD4^BD2^+VHL, and the origin data was listed in Figure 4A to analyze further. Similarly, Benjamin L. Ebert’s group^27^ using ITC systematically detected the half circle of molecular glue CR8 with DDB1 and CDK12-cycK, and determined thermodynamic signatures were summarized in Figure 4C.

**Figure 3.**
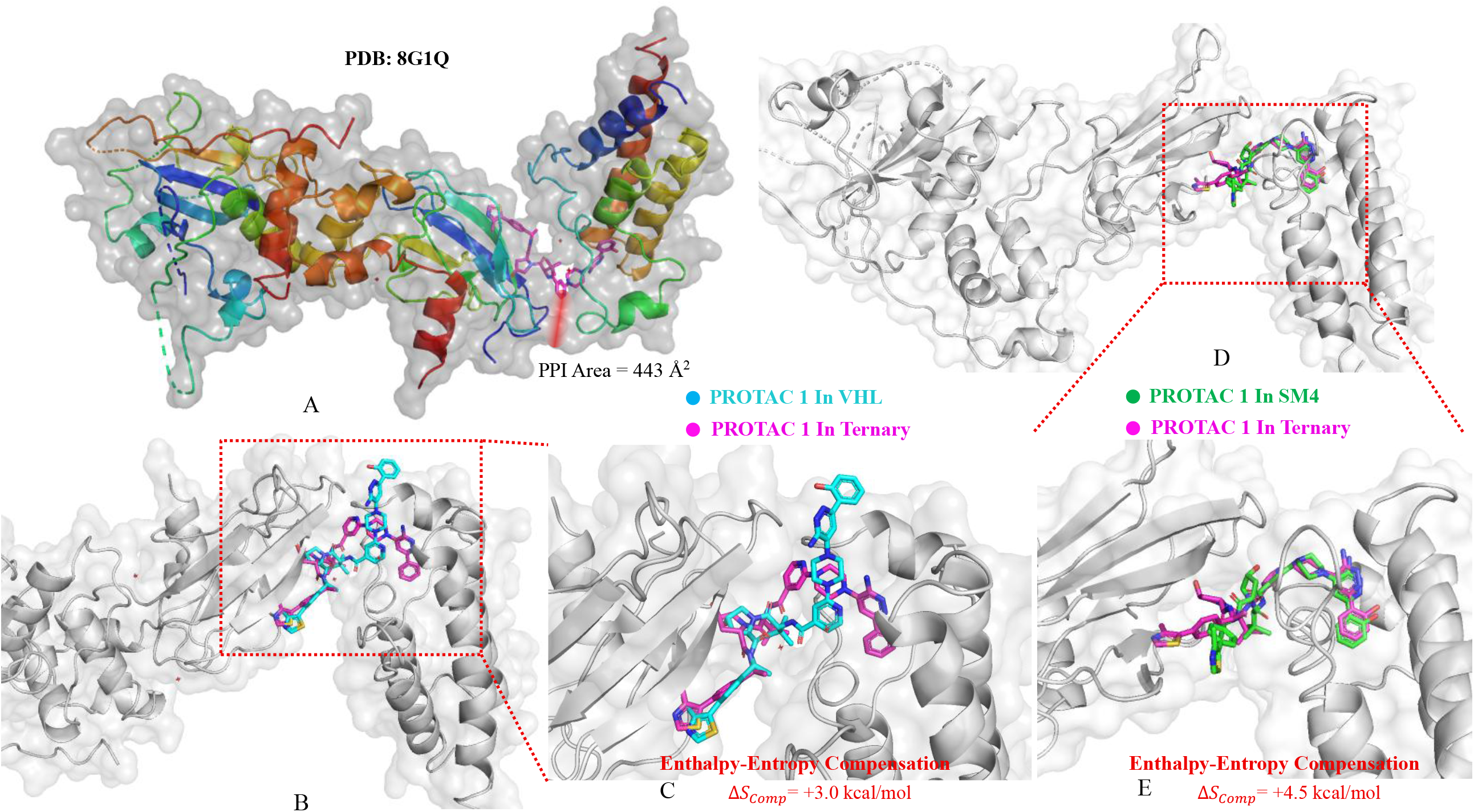
Binding poses analysis of ternary complex SM4-1-VBS (PDB:8G1Q). A) PPI interface of ternary complex. B)The binding pose of compound 1 in VBC (blue) and Ternary complex (pink). C) The enlarged part of image B. D) The binding pose of compound 1 in SM4 (Green) and Ternary complex (pink). C) The enlarged part of image D.

**Figure 4.**
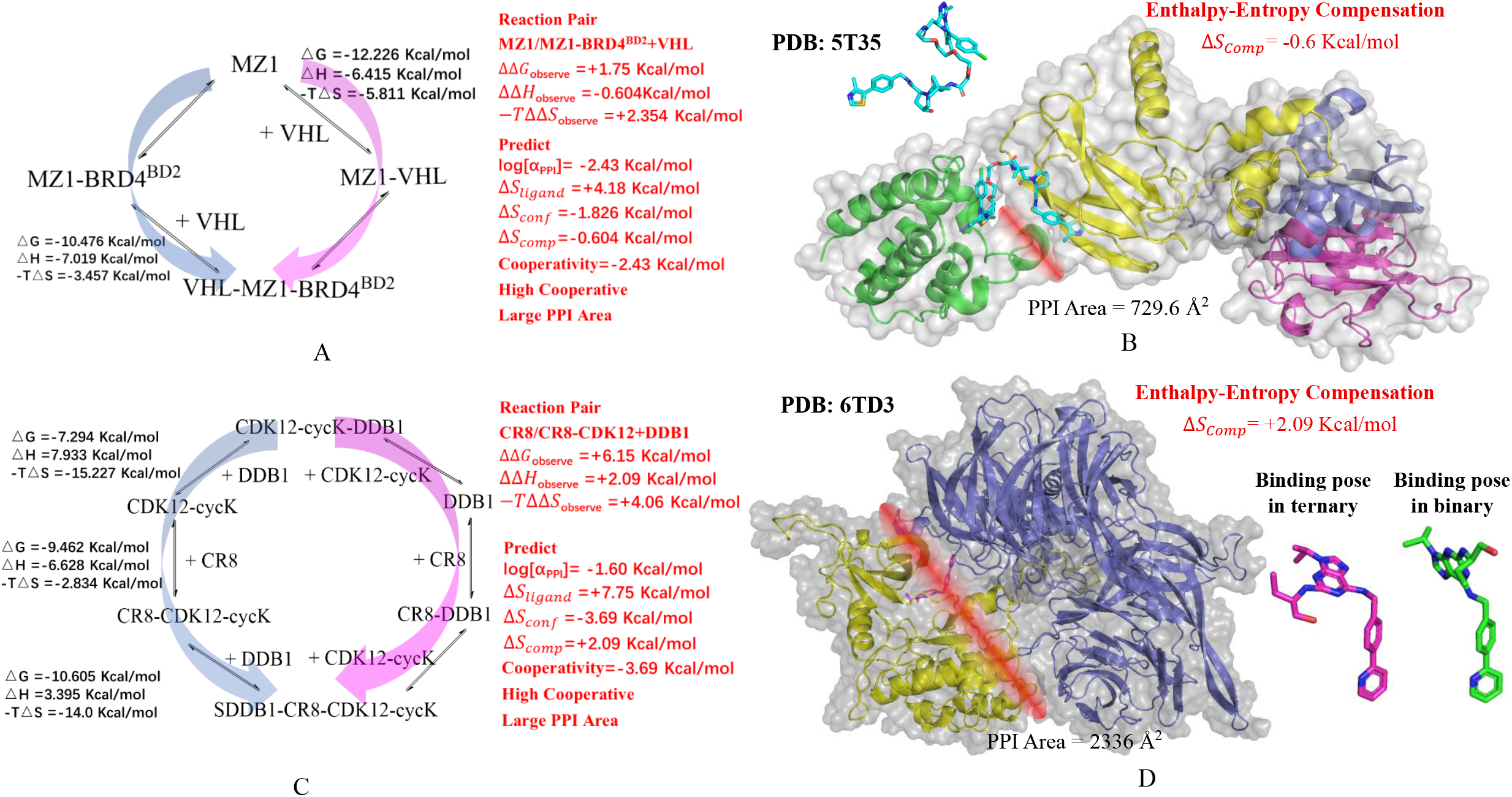
A) Thermodynamic titration Data of reaction pair of MZ1/MZ1-BRD4^BD2^+VHL. B) The PPI buried area of VHL-MZ1-BRD4^BD2^ ternary complex and the pose of the PROTAC. A) Thermodynamic titration Data of reaction pair of CR8/CR8-CDK12+DDB1. B) The PPI buried area of DDB1-CR8-CDK12-cycK ternary complex and the pose of the PROTAC in ternary complex (left of right corner) and binary complex (right of right corner),respectively.

## DISCUSSION

The decade-long confusion surrounding cooperativity in PROTAC and molecular-glue ternary complexes has now been resolved. Both the classical (Douglass) definition and the Ciulli definition of α are thermodynamically correct, yet they quantify entirely different physical effects. The classical α ≫ 1 that dominates the literature does not reflect steric clash or negative cooperativity at the induced POI–E3 interface. Instead, it almost entirely arises from a universal, mass-dependent entropic penalty paid when a ∼1 kDa bifunctional molecule is converted into a 40–200 kDa “gigantic ligand” upon forming the first binary complex. This Δ*S*_*ligand*_ term, typically +4∼+8 kcal·mol^−1^ for PROTAC, is unavoidable in any induced-proximity system and quantitatively explains why Douglass’ α ≫ 1 is observed in 100 % of reported degraders, even when cryo-EM structures reveal extensive, favorable POI– E3 contacts.

Ciulli’s α, by contrast, is the composite of this gigantic-ligand entropy penalty and the genuine protein–protein cooperativity log[*α*]_*PPI*_ that medicinal chemists intuitively wish to maximize. Because Δ*S*_*ligand*_ is large and unfavorable, true positive cooperativity at the induced interface is frequently masked in the modest 10∼1000-fold range (log[*α*]_*PPI*_≈1∼4), yet this is more than sufficient to drive robust degradation when superimposed on high-affinity warheads. Our thermodynamic dissection therefore reconciles the empirical success of Ciulli’s α as a design parameter with the absence of the catastrophic steric penalties that classical α ≫ 1 had seemed to imply.

### Host Force Field “Full-embedded” Model Masked Ligand Entropy

Sackur-Tetrode equation of translational entropy was rigorously deducted since 1912. Finkelstein/Janin(1989), Page/Jencks(1971) and others did calculate translational/rotational entropy also rigorously.^28-31^ All works were clearly showed the relationship between translational entropy and mass of the ligand as *S*_*trans*_ ∝ *In*M. However, in experiment datasets, translational entropy penalty looked like a constant, not a variable one worth emphasizing. Thus, in traditional dynamic model, translational entropy was usually regarded as a slight fluctuating constant correction (∼1–2 kcal/mol). And this treatment was perfectly fit the experiment results. So, why experiment datasets didn’t show this mass-dependent translational entropy?

In conventional small-molecule binding, ligands are fully embedded within the host protein’s force field.^32-35^ The protein’s dense force field was just like a “black box”. As molecular weight of ligand increases, its buried surface area typically scales accordingly, providing additional favorable enthalpic interactions (van der Waals, hydrophobic burial, H-bonds, or dispersing water).^36,37^ This “force density” within the pocket ensures enthalpy gains compensate the rising translational/rotational entropy penalty, resulting in classic enthalpy/entropy or entropy/entropy compensation. These compensation masks the penalty in observable ΔG, rendering it undetectable in classical systems. Consequently, binding affinity assay shows little systematic mass-dependence. FEP implementations apply a constant crate/standard-state entropy term to account for translational/rotational loss. This assumes were based on similar-sized partners and full embedding.^38-42^ Similar strategies were applied in other models. For MM-GBSA/PBSA, entropy is estimated via normal-mode analysis or quasi-harmonic, good for conformational entropy but poor for absolute translational/rotational entropy.^43,44^ Docking Scores were empirical but no explicit mass-dependent entropy.^45-47^ These methods were excellent performance in fully embedded, compensable host force field but developed and benchmarked on classical drug-like datasets.

In partial-buried/asymmetric systems (PROTAC, binary complexes), the “giant” exposed part adds mass without proportional new favorable contacts (low force density outside the pocket), no compensatory enthalpy from extra surface area for the exposed bulk, thus the entropy penalty is uncompensated leading to large unfavorable ΔG contribution, visible as massive affinity drops or apparent negative Cooperativity. This is why classic drug discovery (fully embedded ligands) could safely ignore mass-dependent entropy: nature evolution and chemists design ensured compensation via optimized interfaces. But induced-proximity breaks that assumption by design, the linker/binary is deliberately non-interacting/exposed. In a word, induced-proximity systems feature partial-buried “giant ligands” with substantial solvent-exposed mass contributing minimal compensatory interactions, unmasking the full entropy penalty.

To validate the ligand entropy calculation function, we analyzed 9,380 PROTAC records in the PROTAC-DB database, the IC_50_ records of many examples in the database confirm that the mass dependent rules especially when linker part was gradually increased (Figure S3, S4). We further use formula (4) to calculate the IC_50_ value from literature and found calculated results were matched the report result well within ∼20% error. Apparently, the results from the function (4) were much more precise than traditional commercial software (Table S4). Impressively, the tendency trend of these predicted IC_50_ value by commercial software was reversed to reported IC_50_ revealing that this traditional model was not suitable for PROTAC.

### ITC Thermodynamic Cycle Data Confirmed Giant Ligand Entropy

A central tenet of our proposed model is that progression from a binary complex to a ternary complex entails a substantial increase in molecular weight, resulting in a remarkable reduction of extra ligand entropy Δ*S*_*ligand*_. For instance, using an empirical approximation derived from equation (4), recruitment of a ∼40 kDa protein to a PROTAC-bound binary complex (∼1000 Da PROTAC) incurs an entropic penalty of approximately +5.7 kcal/mol, corresponding to a ∼50,000-fold attenuation in binding affinity (for example, *K*_*d*_ from 10 nM increasing to 500 *μ*M). Thus, observation of an anomalously large unfavorable entropic contribution in the binary-to-ternary transition would strongly support our ligand entropy-penalty theory.

Analysis of the thermodynamic cycle proceeding clockwise: Compound 1 → 1-VBC → SM2-1-VBC, we noticed that VBC part is huge with molecular weight 41.226 kDa, significantly larger than the SM2 or SM4 bromodomains (∼14.4 and ∼14.7 kDa). Consequently, formation of the ternary complex from the preformed 1·VBC binary requires immobilization of the much larger “giant ligand” (1-VBC), incurring a substantial ligand entropic penalty (Δ*S*_*ligand*_) upon restriction of its translational and rotational degrees of freedom. Our model predicts this ligand entropy penalty (Δ*S*_*ligand*_) to be approximately +6 kcal/mol, which is enough to weaken binding affinity by ∼100,000-fold. Experimental ITC data closely match this prediction, showing unfavorable entropic contributions (–TΔΔS) of +5.4 kcal/mol and +5.7 kcal/mol for the SM2 and SM4 systems, respectively (Figure 2). This massive entropy loss cannot be explained by normal thermodynamic models. Even whole changed enthalpy with +3.1 kcal/mol and +3.0 kcal/mol, respectively, still can’t compensate for this massive entropy loss. An explanation was “thermodynamic cooperativity,” but the direct protein–protein interface between SM4 and VBC is ridiculously small — only 443 Å^2^, just 3.2% of the SM4 surface (Figure 3A). That’s tiny PPI surface is incredibly paying a monstrous entropy price driving the ternary complex.

Consistent patterns emerge from additional ITC datasets. Holly Soutter’s group titrates the reaction pair of MZ1/MZ1-BRD4^BD2^+VHL, and the result showed unfavorable entropic term of approximately +2.4 kcal/mol in the relevant cycle arm (Figure 4A). Similarly, systematic ITC evaluation of the molecular glue CR8 with the DDB1-CDK12-cyclin K system yielded an observed giant entropic penalty of +4.06 kcal/mol (Figure 4C). These findings across diverse systems underscore that ternary complexes often exhibit unfavorable thermodynamics dominated by this huge ligand-entropy penalty rather than Cooperativity at the induced protein–protein interface.

### True Entropic Positive Cooperativity and Enthalpy-Entropy Compensation

In induced-proximity systems such as ternary complex-forming degraders, true thermodynamic cooperativity arises predominantly from an entropic contribution associated with the restriction of conformational flexibility (side-chain and backbone fluctuations) upon formation of the induced protein–protein interface (POI–E_3_) and within intramolecular domains of the complex. Consequently, the discrepancy observed between the computationally predicted ligand entropy penalty (Δ*S*_*ligand*_) and the experimentally measured total entropy (ΔΔ*S*_observed_) can be attributed to this additional cooperativity at the PPI interface, which manifests as the conformational entropy Δ*S*_*Conf*_ of the entire ternary complex. From ITC thermodynamic parameters, the conformational entropy Δ*S*_*Conf*_ was calculated to be –0.572 kcal/mol for the SM2-1-VBC ternary complex and –0.272 kcal/mol for the SM4-1-VBC ternary complex. The substantially more negative Δ*S*_*Conf*_ value for SM2 indicates stronger positive Cooperativity in the SM2/VBC interaction compared with SM4/VBC.

Experimentally derived total ΔΔG values were +8.5 kcal/mol for 1/1-VBC+SM2 reaction pair and +8.7 kcal/mol for 1/1-VBC+SM4. Subtracting entropic terms from the total ΔΔG, it revealed unfavorable enthalpic contributions of approximately +3.1 kcal/mol (SM2) and +3.0 kcal/mol (SM4), which was predicted as compensation entropy penalties from enthalpy in these systems. Because in general, the portion of linker and ligand exposed in solution is unlikely just applying free state poses to perfectly match the binding pose in target protein. Further structural analysis of the binding poses of PROTAC compound 1 in the binary complex 1-VBC versus the ternary complex SM4-1-VBC (Figure 3B,3C) showed substantial differences, particularly in the orientation of the SM4 portion (Figure 3D,3E). This observation indicates that compound 1 is forced into a higher-energy conformation in the ternary complex to satisfy the geometric requirements of both the SM4 and E_3_ ligase (VBC) simultaneously.

Consequently, the observed enthalpic penalty does not arise from unfavorable protein–protein contacts perse, but rather from the internal strain energy stored in the distorted PROTAC ligand upon ternary-complex formation.

### Buried PPI Area was the Key for High Cooperativity

From ITC thermodynamic parameters, the conformational entropy Δ*S*_*Conf*_ was calculated to be –0.572 kcal/mol for the 1-SM2-VBC ternary complex and –0.272 kcal/mol for the 1-SM4-VBC ternary complex. The substantially more negative Δ*S*_*Conf*_ value for SM2 indicates stronger positive Cooperativity in the SM2/VBC interaction compared with SM4/VBC. This observation is consistent with the larger buried surface area at the induced protein-protein interface for SM2/VHL (∼500 Å^2^) relative to SM4/VBC (443 Å^2^), which imposes greater restriction of conformational degrees of freedom and thereby yields a larger favorable entropic contribution to overall binding Cooperativity. For reaction pair of MZ1 /MZ1-BRD4^BD2^+VHL, its conformation entropy Δ*S*_*Conf*_ = −1.826 kcal/mol means high Cooperativity and potential larger PPI area. Actually, the PPI area was confirmed by Cryo-EM and be calculated as 729.6 Å^2^, which was larger than SM2/SM4 cases(Figure 4B). These analysis were supporting high Cooperativity related with its buried interface area. Furthermore, for DDB1-CR8-CDK12-cycK complex, its conformation entropy was much larger (-3.69 kcal/mol). Since PPI area of DDB1-CR8-CDK12-cycK complex (2336 Å^2^) was the largest PPI area among known PROTAC and molecular glue (Figure 4D), the true Cooperativity of PROTAC and molecular glue should be ranged in 0∼−4 kcal/mol. It revealed that the buried PPI area was the key for high Cooperativity.

### Flexible Ligand Pays More Enthalpy Penalty in Binding Process

We analyzed the entropic compensation in thermodynamic cycles involving different reaction pairs. For the reaction pairs 1/1-VBC+SM4 (clockwise) and 1/1-SM4+VBC(Anti-clockwise), the entropic compensation terms were +3.0 kcal/mol and +4.5 kcal/mol, respectively (Figure 2). This indicates that when the VHL ligand moiety is solvent-exposed in the free state, formation of the ternary complex incurs a greater entropic penalty than when the SM4 ligand moiety is exposed (Figures 3C and 3E). A similar trend was observed with SM2: the entropic compensation terms for the pairs 1/1-VBC+SM2 and 1/1-SM2+VBC were +3.1 kcal/mol and +4.64 kcal/mol, respectively. Interpreting ternary complex formation as a host–guest binding process provides a straightforward explanation. The SM2/SM4 ligands part of compound 1 are significantly more rigid than the VHL ligand warhead. Rigid ligands possess pre-restricted translational and rotational degrees of freedom in solution, resulting in a smaller loss of translation entropy compared to flexible ligands. Consequently, the system incurs a lower entropic penalty when immobilizing a rigid ligand relative to a flexible one. This principle is well-established in traditional small-molecule drug design, where rigidification of flexible ligands often enhances binding affinity and reduces dissociation constants, typically by 10-to 100-fold, primarily through favorable entropic contributions.

### Less Compensation Entropy of Flexible Linker Enhancing Cooperativity

Compounds 1 was incorporated with rigid linkers, exhibited a substantial unfavorable observed enthalpy change (ΔΔ*H*_observed_>3.0 kcal/mol). This penalty arises because the linker and ligand portions exposed to solvent in their free state are unlikely to adopt conformations that precisely match the bound pose within the target protein. When a PROTAC bearing a rigid linker is required to form a ternary complex, it must adopt a higher-energy conformation to simultaneously satisfy the geometric constraints imposed by both the protein of interest (POI) and the E3 ligase. In contrast, the flexible polyethylene glycol(PEG) linker-based PROTAC MZ1 displayed a favorable enthalpy gain (ΔΔ*H*_observed_=–0.6 kcal/mol).(Figure 4B) This indicates that, during ternary complex formation, the PEG-based flexible linker incurs no conformational strain penalty and instead contributes additional binding enthalpy. This enhancement likely results from favorable interactions between the linker and the surfaces of the POI and E3 ligase. Consequently, flexible linkers appear more effective to promoting higher Cooperativity in ternary complex formation.

Benjamin L. Ebert’s group using ITC systematically detected the half circle of molecular glue CR8 with DDB1 and CDK12-cycK, the results were still aligned with our theory well. The dramatic observed entropy loss was +4.06 kcal/mol. When subtracting the predicted ligand entropy, the real Cooperativity was -3.69 kcal/mol, which also means favorite binding. We noticed that the compensation entropy was +2.09 kcal/mol, which means CR8 should adjust the conformation to fit the best neo-protein interface. The Cryo-EM result shows slightly distorter of the conformation in ternary complex than its in binary complex (Figure 4D). This ITC data further confirmed that rigid linker paying more unfavorable compensation entropy penalty than flexible linker.

### Enlightenment for Rational Design of PROTAC

So, rigid linkers reduce the conformational entropy cost of the free ligand but dramatically increase the enthalpic strain required to adopt the precise geometry demanded by the ternary complex. Flexible linkers pay a higher entropic price in the unbound state but incur almost no strain upon complexation, often yielding higher net cooperativity.

The findings provide insights into the rational design of orally bioavailable PROTAC degraders. During the absorption process, PROTAC incorporating flexible ligands or linkers incur a greater entropic penalty upon conformational restriction at the intestinal membrane or transporter interface, thereby reducing the likelihood of successful absorption and consequently diminishing overall oral bioavailability. Furthermore, flexible linkers may enhance positive Cooperativity in ternary complex formation of POI, but this often decreases binding energetics for unfavorable off-target proteins or receptors, increased off-target interactions, and elevated risks of adverse effects and toxicity.

In contrast, rigid PROTACs, being pre-organized in a conformation approximating the bound state, require less entropic cost for initial capture and absorption, suggesting improved absorption potential and higher oral bioavailability. Upon formation of the ternary complex (POI–PROTAC–E_3_ ligase), a rigid linker must compensate by paying a greater entropic penalty to achieve the precise geometric requirements of the productive complex. This constraint limits excessive Cooperativity, potentially resulting in fewer off-target engagements and reduced toxicity compared to flexible analogs.

Based on these thermodynamic and entropic considerations, rigid linkers appear superior to flexible linkers with respect to both oral bioavailability and safety profile in PROTAC design.

### “Ligand Entropy Barrier Wall/Favorable Cooperativity(*α*) Ladder” Pair Serves Thermodynamic and Kinetic “Off-Switch” for Toxicity

Classical ligand-binding thermodynamics shows that for a typical small-molecule ligand (∼500 Da), this unfavorable contribution is already substantial.^48^ When the “ligand” is instead a pre-assembled PROTAC binary complex (∼50-200 kDa), the ligand entropy penalty often reaches to +4∼+8 kcal/mol. For a polymer/antibody/nanoparticle conjugate (10^3^–10^7^ kDa), the penalty rises dramatically, often reaching +10 to +15 kcal/mol or even higher^49-53^. This gigantic entropic barrier must be compensated by favorable interaction energy for binding to occur at biologically relevant concentrations.^54,55^ Off-target proteins rarely possess interfaces capable of delivering such enormous compensatory energy. Consequently, binding of the giant conjugate is thermodynamically prohibitive, even if the conjugated toxin itself has picomolar binary affinity. In contrast, the intended target site (e.g., a tumor cell overexpressing HER2, folate receptor-α, or a neoepitope recognized by a targeting ligand) can engage in high-avidity or high-Cooperativity interactions that provide the necessary energetic payback, thereby overcoming the translational/rotational entropy wall like a ladder. This principle explains the remarkably wide therapeutic windows of approved antibody–drug conjugates (ADCs), polymer–prodrugs, and nanoparticle-delivered cytotoxins, despite payloads with free-molecule IC_50_ values in the picomolar to femtomolar range.^56,57^ In extreme cases (e.g., >50 nm nanoparticles), the penalty becomes so large, and steric constraints so severe, that off-target binding is not merely unfavorable but physically impossible under physiological conditions. The “ligand entropy barrier wall/favorable Cooperativity(*α*) ladder” pair constituting a true thermodynamic and kinetic “off switch” for toxicity.

## CONCLUSION

In summary, Cooperativity in induced-proximity pharmacology is governed by two energetically comparable terms including induced PPI cooperativity (mostly entropic), and gigantic-ligand entropy penalty, rather than by a single mysterious α value. Seprerating the ligand entropy form the system, two conflicting Cooperativity definitions was reconciled, yielding true path-independent positive PPI Cooperativity. ITC data showed rigid linkers appear superior to flexible linkers with respect to both oral bioavailability and safety profile in PROTAC design. “ligand entropy barrier wall/favorable Cooperativity α ladder” pair is not only impact PROTAC or Molecular Glue system but also constitute the physical basis for all biosystems.

## SUPPORTING INFORMATION

The authors confirm that Supplementary Tables and Figures were available within the supplementary materials.

## ACKNOWLEDGEMENTS

The authors gratefully acknowledge the financial aid from his tutor professor Yan He in Tsinghua University and the opening fund of The State Key Laboratory of Bioorganic Phosphorus Chemistry & Chemical Biology in Tsinghua University. The authors also gratefully appreciate the financial aid from the opening fund of the State Key Laboratory of Chemo/Biosensing and Chemometrics in Hunan University. The author gratefully thanks for the opportunities to improve myself at University of California, Riverside.

The authors thanks the developers for the openly accessible database [PROTAC-DB](http://cadd.zju.edu.cn/protacdb/about) 3.0 and [MolGlueDB](http://cadd.zju.edu.cn/protacdb/about).

## ETHICAL APPROVAL

The authors declare that there is no ethical risk, and the experiment was not involving any animals and no human participants in this work.

## DECLARATION ON THE USE OF GENERATIVE AI

We gratefully acknowledge Grok (xAI) for valuable assistance in English language polishing, structural organization, and insightful scientific discussions that helped refine the concepts presented in this work. The corresponding author Xuan Cao takes full responsibility for the content of the publication.

## TOC

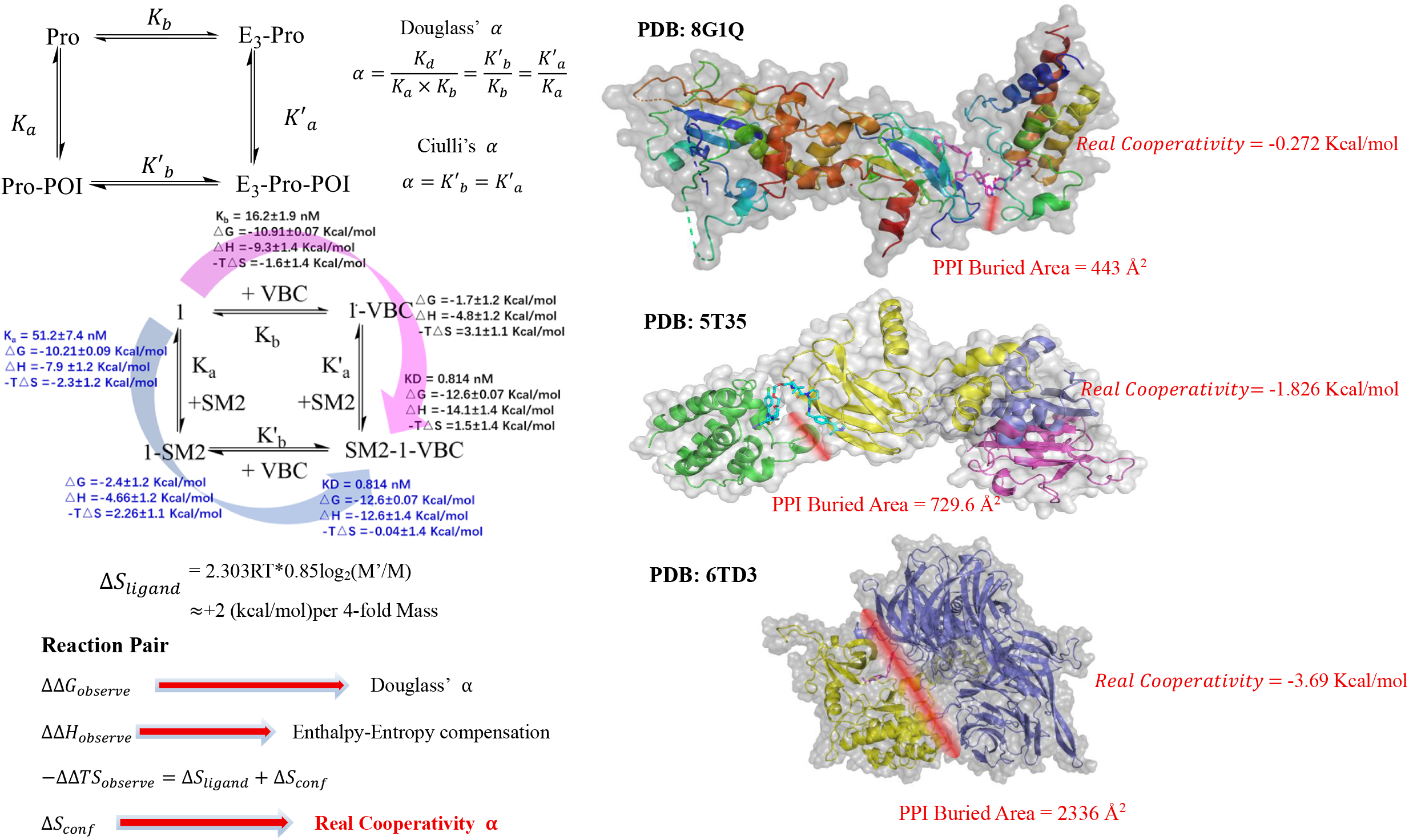

## REFERENCES

1 Mayor-Ruiz, C. et al. Rational discovery of molecular glue degraders via scalable chemical profiling. Nat Chem Biol 16, 1199–1207, doi:10.1038/s41589-020-0594-x (2020).

2 Hassaan, E. Molecular glue modulates mitochondria. Nat Chem Biol 20, 1388, doi:10.1038/s41589-024-01769-0 (2024).

3 Li, J. et al. Activation of human STING by a molecular glue-like compound. Nat Chem Biol 20, 365–372, doi:10.1038/s41589-023-01434-y (2024).

4 Preti, D., Albanese, V. & Marconi, P. C. R. PROTAC-ing tuberculosis. Nat Chem Biol 20, 668–670, doi:10.1038/s41589-024-01624-2 (2024).

5 Crunkhorn, S. PROTAC for p300/CBP in prostate cancer. Nat Rev Drug Discov 24, 906, doi:10.1038/d41573-025-00177-w (2025).

6 Neven, P. & Han, S. N. PROTAC SERD vepdegestrant outperforms fulvestrant for advanced-stage ER(+)HER2(-) breast cancer harbouring acquired ESR1 mutations. Nat Rev Clin Oncol 22, 709–710, doi:10.1038/s41571-025-01062-6 (2025).

7 Alkhalaf, L. M. et al. Thoughts for the future. Nat Chem Biol 21, 6–15, doi:10.1038/s41589-024-01802-2 (2025).

8 Cao, Y., Harris, A. L. & Ciulli, A. Branching beyond bifunctional linkers: synthesis of macrocyclic and trivalent PROTACs. Nat Protoc, doi:10.1038/s41596-025-01283-0 (2025).

9 Dikic, I. et al. Opportunities in proximity modulation: Bridging academia and industry. Mol Cell 85, 3012–3022, doi:10.1016/j.molcel.2025.07.018 (2025).

10 Hinterndorfer, M., Spiteri, V. A., Ciulli, A. & Winter, G. E. Targeted protein degradation for cancer therapy. Nat Rev Cancer 25, 493–516, doi:10.1038/s41568-025-00817-8 (2025).

11 Hughes, S. J. et al. Mode of action of a DCAF16-recruiting targeted glue that can selectively degrade BRD9. Nat Commun 16, 8516, doi:10.1038/s41467-025-63594-w (2025).

12 Lloyd, H. C. et al. A method for the detection and enrichment of endogenous cereblon substrates. Cell Chem Biol 32, 1028–1041 e1013, doi:10.1016/j.chembiol.2025.07.002 (2025).

13 Ma, N. et al. Frustration in the protein-protein interface plays a central role in the cooperativity of PROTAC ternary complexes. Nat Commun 16, 8595, doi:10.1038/s41467-025-63713-7 (2025).

14 Douglass, E. F., Jr., Miller, C. J., Sparer, G., Shapiro, H. & Spiegel, D. A. A comprehensive mathematical model for three-body binding equilibria. J Am Chem Soc 135, 6092–6099, doi:10.1021/ja311795d (2013).

15 Gadd, M. S. et al. Structural basis of PROTAC cooperative recognition for selective protein degradation. Nat Chem Biol 13, 514–521, doi:10.1038/nchembio.2329 (2017).

16 Roy, M. J. et al. SPR-Measured Dissociation Kinetics of PROTAC Ternary Complexes Influence Target Degradation Rate. ACS Chem Biol 14, 361–368, doi:10.1021/acschembio.9b00092 (2019).

17 Diehl, C. J., Salerno, A. & Ciulli, A. Ternary Complex-Templated Dynamic Combinatorial Chemistry for the Selection and Identification of Homo-PROTACs. Angew Chem Int Ed Engl 63, e202319456, doi:10.1002/anie.202319456 (2024).

18 Popow, J. et al. Targeting cancer with small-molecule pan-KRAS degraders. Science 385, 1338–1347, doi:10.1126/science.adm8684 (2024).

19 Wurz, R. P. et al. Affinity and cooperativity modulate ternary complex formation to drive targeted protein degradation. Nat Commun 14, 4177, doi:10.1038/s41467-023-39904-5 (2023).

20 Konstantinidou, M. & Arkin, M. R. Molecular glues for protein-protein interactions: Progressing toward a new dream. Cell Chem Biol 31, 1064–1088, doi:10.1016/j.chembiol.2024.04.002 (2024).

21 Geiger, T. M. et al. Discovery of a Potent Proteolysis Targeting Chimera Enables Targeting the Scaffolding Functions of FK506-Binding Protein 51 (FKBP51). Angew Chem Int Ed Engl 63, e202309706, doi:10.1002/anie.202309706 (2024).

22 Vetma, V. et al. Identification of a Highly Cooperative PROTAC Degrader Targeting GTP-Loaded KRAS(On) Alleles. J Am Chem Soc 147, 41367–41378, doi:10.1021/jacs.5c10354 (2025).

23 Grimus, W. Statistische Physik und Thermodynamik: Grundlagen und Anwendungen. 2., überarbeitete Auflage. edn, (De Gruyter, 2015).

24 Finkelstein, A. V. & Janin, J. The price of lost freedom: entropy of bimolecular complex formation. Protein Eng 3, 1–3, doi:10.1093/protein/3.1.1 (1989).

25 Shcherbakov, A. A., Poppe, L. & Vaish, A. Biophysical Approaches for Investigating the Dynamics and Cooperativity of Ternary Complexes in Targeted Protein Degradation. J Med Chem 68, 12904–12910, doi:10.1021/acs.jmedchem.5c00652 (2025).

26 Jiang, W. & Soutter, H. The Development and Application of Biophysical Assays for Evaluating Ternary Complex Formation Induced by Proteolysis Targeting Chimeras (PROTACS). J Vis Exp, doi:10.3791/65718 (2024).

27 Slabicki, M. et al. The CDK inhibitor CR8 acts as a molecular glue degrader that depletes cyclin K. Nature 585, 293–297, doi:10.1038/s41586-020-2374-x (2020).

28 Murphy, K. P., Xie, D., Thompson, K. S., Amzel, L. M. & Freire, E. Entropy in biological binding processes: estimation of translational entropy loss. Proteins 18, 63–67, doi:10.1002/prot.340180108 (1994).

29 Amzel, L. M. Loss of translational entropy in binding, folding, and catalysis. Proteins 28, 144–149 (1997).

30 Siebert, X. & Amzel, L. M. Loss of translational entropy in molecular associations. Proteins 54, 104–115, doi:10.1002/prot.10472 (2004).

31 Piantadosi, S. Translational clinical trials: an entropy-based approach to sample size. Clin Trials 2, 182–192, doi:10.1191/1740774505cn078oa (2005).

32 Yin, J., Fenley, A. T., Henriksen, N. M. & Gilson, M. K. Toward Improved Force-Field Accuracy through Sensitivity Analysis of Host-Guest Binding Thermodynamics. J Phys Chem B 119, 10145–10155, doi:10.1021/acs.jpcb.5b04262 (2015).

33 Bell, D. R. et al. Calculating binding free energies of host-guest systems using the AMOEBA polarizable force field. Phys Chem Chem Phys 18, 30261–30269, doi:10.1039/c6cp02509a (2016).

34 Henriksen, N. M. & Gilson, M. K. Evaluating Force Field Performance in Thermodynamic Calculations of Cyclodextrin Host-Guest Binding: Water Models, Partial Charges, and Host Force Field Parameters. J Chem Theory Comput 13, 4253–4269, doi:10.1021/acs.jctc.7b00359 (2017).

35 Slochower, D. R. et al. Binding Thermodynamics of Host-Guest Systems with SMIRNOFF99Frosst 1.0.5 from the Open Force Field Initiative. J Chem Theory Comput 15, 6225–6242, doi:10.1021/acs.jctc.9b00748 (2019).

36 Amadasi, A. et al. Explaining cyclodextrin-mycotoxin interactions using a ‘natural’ force field. Bioorg Med Chem 15, 4585–4594, doi:10.1016/j.bmc.2007.04.006 (2007).

37 Wang, S. M. et al. Chiral recognition of neutral guests by chiral naphthotubes with a bis-thiourea endo-functionalized cavity. Nat Commun 14, 5645, doi:10.1038/s41467-023-41390-8 (2023).

38 Perthold, J. W., Petrov, D. & Oostenbrink, C. Toward Automated Free Energy Calculation with Accelerated Enveloping Distribution Sampling (A-EDS). J Chem Inf Model 60, 5395–5406, doi:10.1021/acs.jcim.0c00456 (2020).

39 Cappel, D., Mozziconacci, J. C., Braun, T. & Steinbrecher, T. Performance of Relative Binding Free Energy Calculations on an Automatically Generated Dataset of Halogen-Deshalogen Matched Molecular Pairs. J Chem Inf Model 61, 3421–3430, doi:10.1021/acs.jcim.1c00290 (2021).

40 Petrov, D. Perturbation Free-Energy Toolkit: An Automated Alchemical Topology Builder. J Chem Inf Model 61, 4382–4390, doi:10.1021/acs.jcim.1c00428 (2021).

41 Pitman, M., Hahn, D. F., Tresadern, G. & Mobley, D. L. To Design Scalable Free Energy Perturbation Networks, Optimal Is Not Enough. J Chem Inf Model 63, 1776–1793, doi:10.1021/acs.jcim.2c01579 (2023).

42 Chen, L. et al. Performance and Analysis of the Alchemical Transfer Method for Binding-Free-Energy Predictions of Diverse Ligands. J Chem Inf Model 64, 250–264, doi:10.1021/acs.jcim.3c01705 (2024).

43 Belhassan, A. et al. Camphor, Artemisinin and Sumac Phytochemicals as inhibitors against COVID-19: Computational approach. Comput Biol Med 136, 104758, doi:10.1016/j.compbiomed.2021.104758 (2021).

44 Ivanova, A., Mokshyna, O. & Polishchuk, P. StreaMD: the toolkit for high-throughput molecular dynamics simulations. J Cheminform 16, 123, doi:10.1186/s13321-024-00918-w (2024).

45 Erdogan, A. et al. Design, synthesis and evaluation of novel oxadiazole-based compounds as potential Rac1 inhibitors against breast cancer. Eur J Med Chem 303, 118396, doi:10.1016/j.ejmech.2025.118396 (2025).

46 Malankar, G. S. et al. Engineering NIR Probes to Enhance Affinity and Clinical Workflow Compatibility for Prostate Cancer Imaging. Angew Chem Int Ed Engl, e20355, doi:10.1002/anie.202520355 (2025).

47 Zeng, W. et al. Discovery of Imidazo[1,2-b]pyridazine Derivatives as Potent PI3K/mTOR Dual Inhibitors for the Treatment of Pulmonary Fibrosis. J Med Chem, doi:10.1021/acs.jmedchem.5c02587 (2025).

48 Zhou, H. X. & Gilson, M. K. Theory of free energy and entropy in noncovalent binding. Chem Rev 109, 4092–4107, doi:10.1021/cr800551w (2009).

49 Krishnamurthy, V. & Chung, S. H. Adaptive Brownian dynamics simulation for estimating potential mean force in ion channel permeation. IEEE Trans Nanobioscience 5, 126–138, doi:10.1109/tnb.2006.875035 (2006).

50 Krishnamurthy, V. M. et al. Thermodynamic parameters for the association of fluorinated benzenesulfonamides with bovine carbonic anhydrase II. Chem Asian J 2, 94–105, doi:10.1002/asia.200600360 (2007).

51 Krishnamurthy, V. M., Semetey, V., Bracher, P. J., Shen, N. & Whitesides, G. M. Dependence of effective molarity on linker length for an intramolecular protein-ligand system. J Am Chem Soc 129, 1312–1320, doi:10.1021/ja066780e (2007).

52 Krishnamurthy, V. V. et al. Orientational distributions and nematic order of rodlike magnetic nanoparticles in dispersions. Phys Rev E Stat Nonlin Soft Matter Phys 77, 031403, doi:10.1103/PhysRevE.77.031403 (2008).

53 Krishnamurthy, V. M. et al. Ligand-induced protein mobility in complexes of carbonic anhydrase II and benzenesulfonamides with oligoglycine chains. PloS one 8, e57629, doi:10.1371/journal.pone.0057629 (2013).

54 Zhang, Y., Li, L. & Wang, J. Tuning cellular uptake of nanoparticles via ligand density: Contribution of configurational entropy. Phys Rev E 104, 054405, doi:10.1103/PhysRevE.104.054405 (2021).

55 Wan, H., Xu, D., Gao, L. & Yan, L. T. Entropy-Mediated Nanoparticle Cellular Uptake. Small Sci 4, 2300078, doi:10.1002/smsc.202300078 (2024).

56 Muro, S. Challenges in design and characterization of ligand-targeted drug delivery systems. J Control Release 164, 125–137, doi:10.1016/j.jconrel.2012.05.052 (2012).

57 Srinivasarao, M. & Low, P. S. Ligand-Targeted Drug Delivery. Chem Rev 117, 12133–12164, doi:10.1021/acs.chemrev.7b00013 (2017).

