## supplemental table and figure for "Reconciling Cooperativity Definition in PROTACs and Molecular Glues: Thermodynamic Dissection into PPI and Ligand Entropic Contributions"

### CAPTION LIST

Table S1. List of calculated Douglass' Cooperativity from database PROTACDB.

Table S2. Statistic PPI of ternary complex of PROTAC

Table S3. List of Ciulli's Cooperativity $\boldsymbol{\alpha}$(POI) and $\boldsymbol{\alpha'}$($E_3$ ligase) from database PROTACDB.

Table S4. The prediction value of $IC_{50}$ or $K'_b$ using the ligand entropy simulation and some commercial software.

Table S5. Reported ITC thermodynamics cycle data of binary/ternary complex from compound 1, VBC , and SM2 /SM4.

Table S6. Revised full ITC thermodynamic data of binary/ternary complex from compound 1, VBC , and SM2 /SM4.

Figure S1. Cryo-EM of Ternary complex structure of PROTAC (PDB:8BB2 and 6W8I) and Ternary complex structure of Molecular Glue (PDB:9NFQ and 909I).

Figure S2. Cases of the report $IC_{50}$ value for the warhead and its related PROTAC molecules.

Figure S3. Cases of the report $IC_{50}$ value of the PROTAC molecules with different length unit's linker.

Table S1. List of calculated Douglass' Cooperativity from database PROTACDB.

| ID | Structure | $K_a$ (nM) | $K_b$ (nM) | $K_d$ (nM) | $K'_b$ (M) | $\alpha$ (Douglass') | Quote from |
|---|---|---|---|---|---|---|---|
| 2892 | 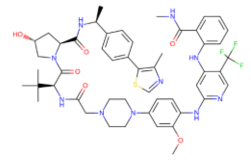 | 1.58 | 1000 | 1.26 | 0.80 | 8.00E+05 | Angew Chem Int Edit. 2021, 60(43): 23327-23334 |
| 2888 | 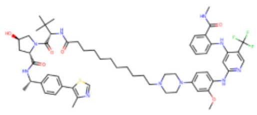 | 7.94 | 1995.3 | 2 | 0.25 | 1.26E+05 | Angew Chem Int Edit. 2021, 60(43): 23327-23334 |
| 2887 | 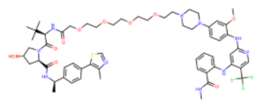 | 0.63 | 316.2 | 5.01 | 7.69 | 2.43E+07 | Angew Chem Int Edit. 2021, 60(43): 23327-23334 |
| 2890 | 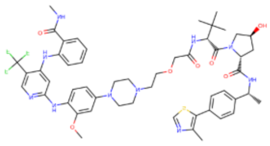 | 1 | 794.3 | 6.31 | 6.25 | 7.87E+06 | Angew Chem Int Edit. 2021, 60(43): 23327-23334 |
| 2889 | 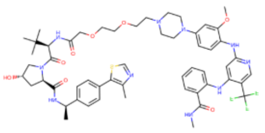 | 1 | 501.2 | 7.94 | 7.69 | 1.53E+07 | Angew Chem Int Edit. 2021, 60(43): 23327-23334 |
| 2891 | 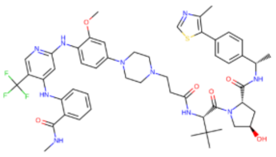 | 1.26 | 1258.9 | 10 | 8.33 | 6.62E+06 | Angew Chem Int Edit. 2021, 60(43): 23327-23334 |
| 797 | 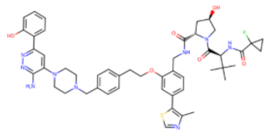 | 550 | 100 | 28 | 0.05 | 5.10E+05 | Nat Chem Biol. 2019, 15(7):672-680 |
| 798 | 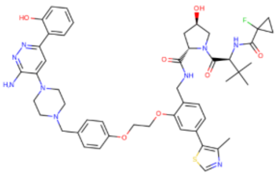 | 504 | 250 | 42 | 0.08 | 3.33E+05 | Nat Chem Biol. 2019, 15(7):672-680 |
| 796 | 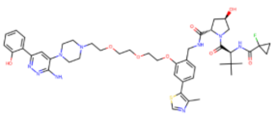 | 1500 | 234 | 220 | 0.15 | 6.28E+05 | Nat Chem Biol. 2019, 15(7):672-680 |
| 1649 | 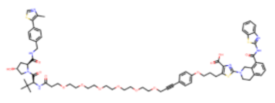 | 0.6 | 190 | 290 | 500.00 | 2.63E+09 | ACS Chem Biol. 2020, 15(9):2316-2323 |
| 1099 | 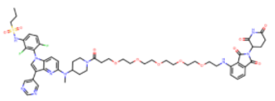 | 25 | 850 | 2170 | 90.91 | 1.07E+08 | Nat Chem Biol. 2020, 16(11):1170-1178 |
| 335 | 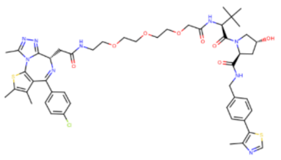 | 62 | 29 | 30 | 0.49 | 1.67E+07 | Nat Chem Biol. 2017, 13(5):514-521 |
| 22 | 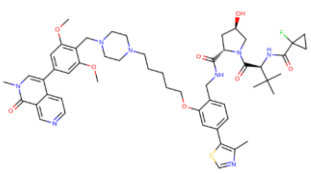 | 5.1 | 26 | 27 | 5.29 | 2.04E+08 | J Med Chem. 2019, 62(2):699-726 |

| ID | Structure | $K_a$ (nM) | $K_b$ (nM) | $K_d$ (nM) | $K'_b$ (M) | $\alpha$ (Douglass') | Quote from |
|---|---|---|---|---|---|---|---|
| 2050 | 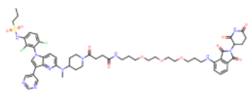 | 82 | 1010 | 2460 | 33.33 | 3.30E+07 | Nat Chem Biol. 2020, 16(11):1170-1178 |
| 1098 | 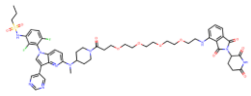 | 12 | 900 | 2570 | 217.39 | 2.42E+08 | Nat Chem Biol. 2020, 16(11):1170-1178 |
| 2049 | 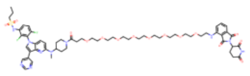 | 85 | 1070 | 2980 | 35.71 | 3.34E+07 | Nat Chem Biol. 2020, 16(11):1170-1178 |
| 2048 | 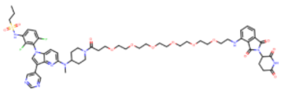 | 83 | 1180 | 3040 | 45.45 | 3.85E+07 | Nat Chem Biol. 2020, 16(11):1170-1178 |
| 1081 | 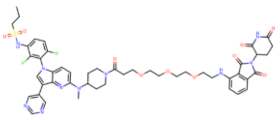 | 65 | 1520 | 3750 | 58.82 | 3.87E+07 | Nat Chem Biol. 2020, 16(11):1170-1178 |
| 796 | 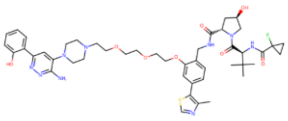 | 2400 SMARCA2 | 234 | 205 | 0.09 | 3.65E+05 | Nat Chem Biol. 2019, 15(7):672-680 |
| 796 | 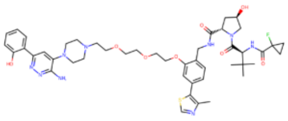 | 3000 SMARCA4 | 234 | 230 | 0.08 | 3.29E+05 | Nat Chem Biol. 2019, 15(7):672-680 |
| 798 | 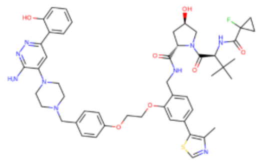 | 770 SMARCA2 | 250 | 26 | 0.03 | 1.35E+05 | Nat Chem Biol. 2019, 15(7):672-680 |
| 798 | 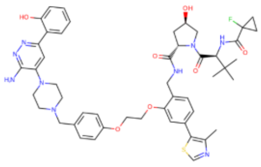 | 560 SMARCA4 | 250 | 51 | 0.09 | 3.67E+05 | Nat Chem Biol. 2019, 15(7):672-680 |
| 797 | 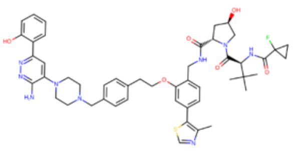 | 790 SMARCA2 | 100 | 45 | 0.06 | 5.71E+05 | Nat Chem Biol. 2019, 15(7):672-680 |
| 1653 | 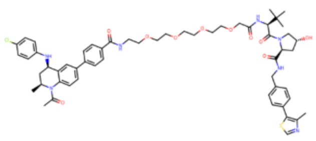 | 45 | 109 | 328 | 7.14 | 6.55E+07 | J Med Chem. 2018, 61(2):504-513 |
| 310 | 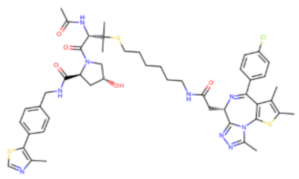 | 75 | 335 | 390 | 5.26 | 1.57E+07 | Nat Chem Biol. 2017, 13(5):514-521 |
| 11 | 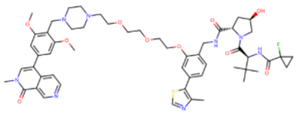 | 8.8 | 87 | 83 | 9.09 | 9.78E+08 | J Med Chem. 2019, 62(2):699-726 |

| ID | Structure | $K_a$ (nM) | $K_b$ (nM) | $K_d$ (nM) | $K'_b$ (M) | $\alpha$ (Douglass') | Quote from |
|---|---|---|---|---|---|---|---|
| 797 | 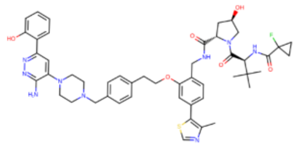 | 990 SMARCA4 | 100 | 76 | 0.08 | 7.68E+05 | Nat Chem Biol. 2019, 15(7):672-680 |
| 1 | 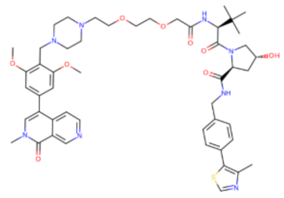 | 15 | 33 | 73 | 5.00 | 1.52E+08 | J Med Chem. 2019, 62(2):699-726 |
| 1653 | 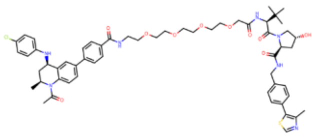 | 39 | 109 | 221 | 5.88 | 5.40E+07 | J Med Chem. 2018, 61(2):504-513 |
| 3150 | 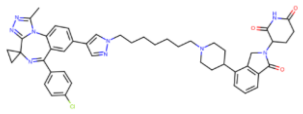 | 58 | 2100 | 4 | 0.07 | 3.28E+04 | ACS Chem Biol. 2021 16(11):2228-2243 |
| 1535 | 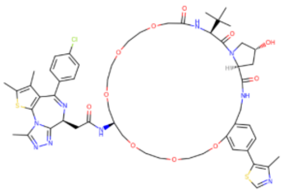 | 180 | 47 | 2 | 0.01 | 2.36E+05 | Angew Chem Int Ed Engl. 2020,59(4):1727-1734 |
| 335 | 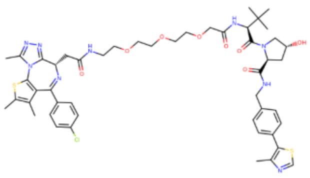 | 26 | 69 | 9 | 0.35 | 5.01E+06 | J Med Chem. 2018, 61(2):504-513 |
| 336 | 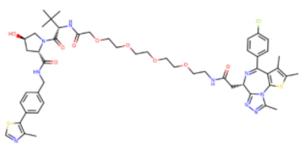 | 27 | 73 | 15 | 0.56 | 7.61E+06 | J Med Chem. 2018, 61(2):504-513 |
| 335 | 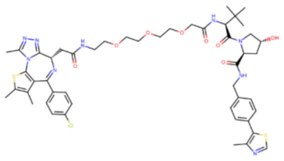 | 62 | 66 | 24 | 0.39 | 5.87E+06 | Nat Chem Biol. 2017, 13(5):514-521 |
| 335 | 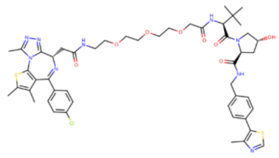 | 39 | 66 | 28 | 0.72 | 1.09E+07 | Nat Chem Biol. 2017, 13(5):514-521 |
| 1652 | 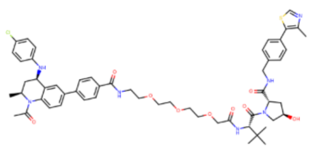 | 4 | 105 | 228 | 58.82 | 5.60E+08 | ACS Chem Biol. 2019, 14(3):361-368 |
| 3254 | 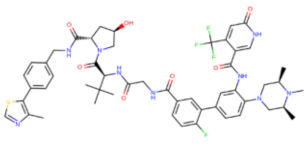 | 122 | 534 | 232 | 1.90 | 3.56E+06 | J Med Chem. 2023 66(23):16168-16186 |
| 185 | 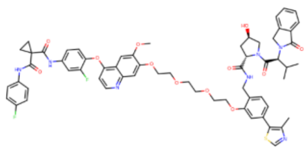 | 550 | 3500 | 436 | 0.79 | 2.27E+05 | Nat Commun. 2019, 10(1):131 |

| ID | Structure | $K_a$ (nM) | $K_b$ (nM) | $K_d$ (nM) | $K'_b$ (M) | $\alpha$ (Douglass') | Quote from |
|---|---|---|---|---|---|---|---|
| 229 | 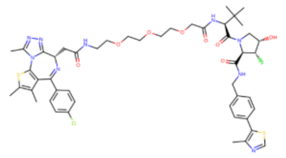 | 28 | 249 | 26 | 0.92 | 3.68E+06 | J Am Chem Soc. 2018, 140(29):9299-9313 |
| 1654 | 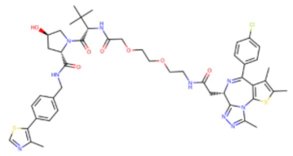 | 17 | 147 | 26 | 1.54 | 1.05E+07 | J Med Chem. 2018, 61(2):504-513 |
| 339 | 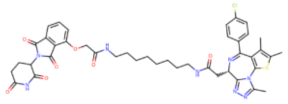 | 46 | 264 | 33 | 0.72 | 2.73E+06 | ACS Chem Biol. 2021, 16(11):2228-2243 |
| 230 | 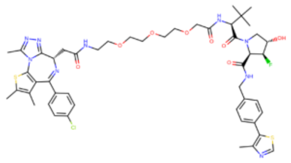 | 32 | 603 | 42 | 1.32 | 2.18E+06 | J Am Chem Soc. 2018, 140(29):9299-9313 |
| 3255 | 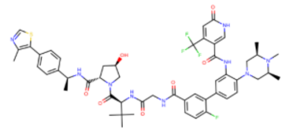 | 63 | 140 | 52 | 0.83 | 5.90E+06 | Sci Transl Med. 2021, 13613:eabj1578 |
| 5634 | 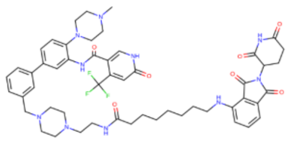 | 125 | 14800 | 63.7 | 0.50 | 3.40E+04 | Oncogene. 2022, 41(24):3328-3340 |
| 229 | 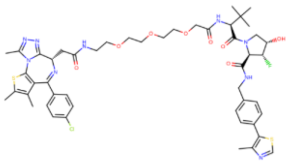 | 97 | 249 | 64 | 0.66 | 2.64E+06 | J Am Chem Soc. 2018, 140(29):9299-9313 |
| 1535 | 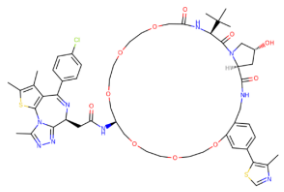 | 743 | 47 | 70 | 0.09 | 2.01E+06 | Angew Chem Int Ed Engl. 2020, 59(4):1727-1734 |
| 2044 | 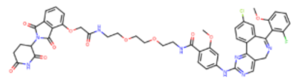 | 375 | 6880 | 183 | 0.49 | 7.09E+04 | Nat Chem Biol. 2020, 16(11):1179-1188 |
| 1652 | 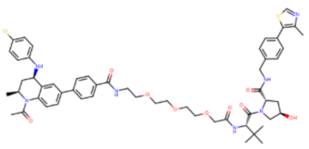 | 4 | 105 | 228 | 58.82 | 5.60E+08 | ACS Chem Biol. 2019, 14(3):361-368 |
| 3254 | 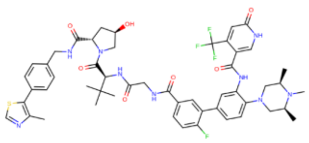 | 122 | 534 | 232 | 1.90 | 3.56E+06 | J Med Chem. 2023 66(23):16168-16186 |
| 185 | 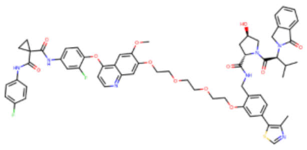 | 550 | 3500 | 436 | 0.79 | 2.27E+05 | Nat Commun. 2019, 10(1):131 |

| ID | Structure | $K_a$ (nM) | $K_b$ (nM) | $K_d$ (nM) | $K'_b$ (M) | $\alpha$ (Douglass') | Quote from |
|---|---|---|---|---|---|---|---|
| 221 | 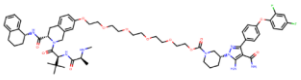 | 259 | 7 | 474 | 1.82 | 2.60E+08 | Nat Chem Biol. 2021, 17(2):152-160 |
| 230 | 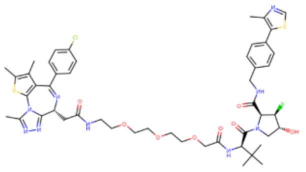 | 111 | 603 | 500 | 4.50 | 7.47E+06 | J Am Chem Soc. 2018, 140(29):9299-9313 |
| 3245 | 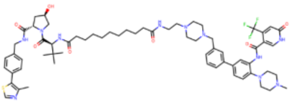 | 120 | 870 | 520 | 4.35 | 5.00E+06 | Sci Transl Med. 2021, 13613:eabj1578 |
| 1651 | 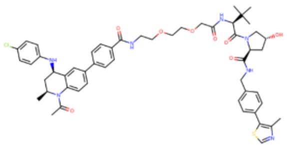 | 3 | 116 | 781 | 250.00 | 2.16E+09 | J Med Chem. 2018, 61(2):504-513 |
| 184 | 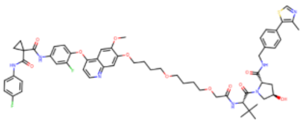 | 500 | 5000 | 1200 | 2.40 | 4.81E+05 | Nat Commun. 2019, 10(1):131 |
| 1653 | 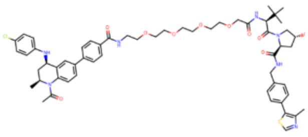 | 39 | 109 | 152 | 3.91 | 3.58E+07 | J Med Chem. 2018, 61(2):504-513 |
| 335 | 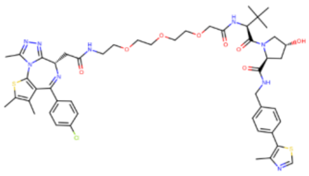 | 21 | 66 | 19 | 0.91 | 1.38E+07 | Nat Chem Biol. 2017, 13(5):514-521 |

Table S2. Statistic PPI of ternary complex of PROTAC.

| PDB entry | POI | E3 ligase | $DC_{50}$ (nM) | Buried PPI Area (Å$^2$) | Buried PPI Area of POI% |
|---|---|---|---|---|---|
| 6w8i | BTK | BIRC2 | 182 | 194.3 | 2.7 |
| 6w7o | BTK | BIRC2 | 800 | 1041.9 | 14.48 |
| 8dso | BTK | BIRC2 | 57 | 637.6 | 8.86 |
| 6boy | BRD4BD1 | CRBN | 6 | 1133.43 | 8.66 |
| 6bnb | BRD4BD1 | CRBN | 500 | 957.22 | 7.31 |
| 6bn9 | BRD4BD1 | CRBN | 5 | 1141.46 | 8.71 |
| 6bn8 | BRD4BD1 | CRBN | 1800 | 1072.71 | 8.19 |
| 6bn7 | BRD4BD1 | CRBN | 50 | 1048.92 | 8.00 |
| 6zhc | BCL-xL | VHL | 4.8 | 669.5 | 0.57 |
| 7pi4 | FAK | VHL | 4 | 739.4 | 1.55 |
| 7khh | BRD4BD1 | VHL | 0.095 | 723.9 | 5.52 |
| 8bds | BRD4BD1 | VHL | 1100 | 624 | 4.77 |
| 8beb | BRD4BD1 | VHL | 320 | 620.3 | 2.44 |
| 6sis | BRD4BD2 | VHL | 25-125 | 680.8 | 0.38 |
| 5t35 | BRD4BD2 | VHL | 8 | 729.6 | 0.41 |
| 7znt | BRD4BD2 | VHL | N.A. | 753.4 | 0.42 |
| 8bdt | BRD4BD2 | VHL | 560 | 724.8 | 0.41 |
| 8bdx | BRD4BD2 | VHL | 1100 | 690 | 0.39 |
| 6hay | SMARCA2 | VHL | 300 | 686.5 | 2.11 |
| 6hax | SMARCA2 | VHL | 70 | 698.3 | 2.15 |
| 7z6l | SMARCA2 | VHL | 78 | 169.9 | 0.52 |
| 7z77 | SMARCA2 | VHL | 2 | 351 | 1.08 |
| 7z76 | SMARCA2 | VHL | 3 | 709.3 | 2.18 |
| 7s4e | SMARCA2 | VHL | 6 | 719.2 | 2.21 |
| 8g1p | SMARCA2 | VHL | 70/221 | 673.8 | 2.07 |
| 6hr2 | SMARCA4 | VHL | 120 | 671.1 | 4.86 |
| 8g1q | SMARCA4 | VHL | N.A. | 443 | 3.21 |
| 7q2j | WDR5 | VHL | 53 | 362.8 | 1.18 |
| 7jtp | WDR5 | VHL | 3.7 | 831.2 | 2.70 |
| 7jto | WDR5 | VHL | 260 | 266.4 | 0.86 |
| 8bb4 | WDR5 | VHL | 116 | 254.3 | 0.83 |
| 8bb2 | WDR5 | VHL | Inactive | 99.7 | 0.32 |
| 8bb5 | WDR5 | VHL | Partial | 181.1 | 0.59 |

Table S3. List of Ciulli's Cooperativity $\alpha$(POI) and $\alpha'$($E_3$ ligase) from database PROTACDB.

| ID | Structure | $K_a$ (nM) | $K_b$ (nM) | $K_d$ (nM) | $\alpha$ (POI) | $\alpha'$ ($E_3$ ligase) | Quote from |
|---|---|---|---|---|---|---|---|
| 2892 | 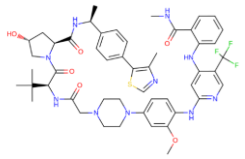 | 1.58 | 1000 | 1.26 | 1.25 | 793 | Angew Chem Int Edit. 2021, 60(43): 23327-23334 |
| 2888 | 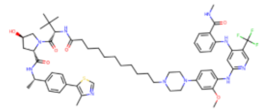 | 7.94 | 1995.3 | 2 | 3.97 | 997.65 | Angew Chem Int Edit. 2021, 60(43): 23327-23334 |
| 2887 | 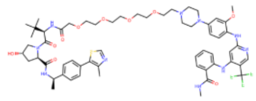 | 0.63 | 316.2 | 5.01 | 0.13 | 63.2 | Angew Chem Int Edit. 2021, 60(43): 23327-23334 |
| 2890 | 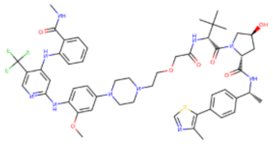 | 1 | 794.3 | 6.31 | 0.16 | 125.8 | Angew Chem Int Edit. 2021, 60(43): 23327-23334 |
| 2889 | 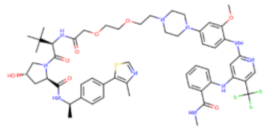 | 1 | 501.2 | 7.94 | 0.13 | 63.1 | Angew Chem Int Edit. 2021, 60(43): 23327-23334 |
| 2891 | 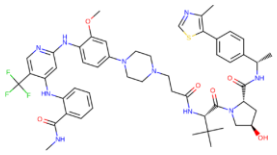 | 1.26 | 1258.9 | 10 | 0.12 | 125.9 | Angew Chem Int Edit. 2021, 60(43): 23327-23334 |
| 797 | 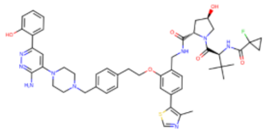 | 550 | 100 | 28 | 19.6 | 3.6 | Nat Chem Biol. 2019, 15(7):672-680 |
| 798 | 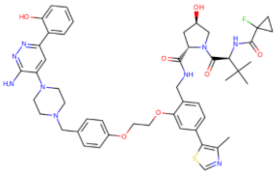 | 504 | 250 | 42 | 12 | 5.95 | Nat Chem Biol. 2019, 15(7):672-680 |
| 796 | 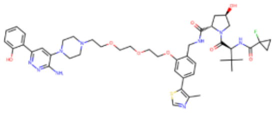 | 1500 | 234 | 220 | 6.8 | 1.06 | Nat Chem Biol. 2019, 15(7):672-680 |
| 1649 | 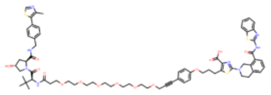 | 0.6 | 190 | 290 | 0.002 | 0.65 | ACS Chem Biol. 2020, 15(9):2316-2323 |
| 1099 | 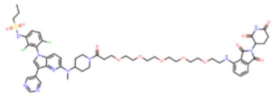 | 25 | 850 | 2170 | 0.011 | 0.39 | Nat Chem Biol. 2020, 16(11):1170-1178 |

| ID | Structure | $K_a$ (nM) | $K_b$ (nM) | $K_d$ (nM) | $\alpha$ (POI) | $\alpha'$ ($E_3$ ligase) | Quote from |
|---|---|---|---|---|---|---|---|
| 2050 | 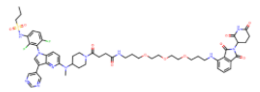 | 82 | 1010 | 2460 | 0.03 | 0.41 | Nat Chem Biol. 2020, 16(11):1170-1178 |
| 1098 | 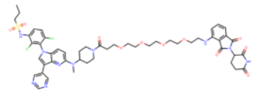 | 12 | 900 | 2570 | 0.0046 | 0.35 | Nat Chem Biol. 2020, 16(11):1170-1178 |
| 2049 | 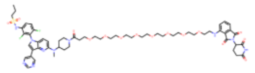 | 85 | 1070 | 2980 | 0.028 | 0.36 | Nat Chem Biol. 2020, 16(11):1170-1178 |
| 2048 | 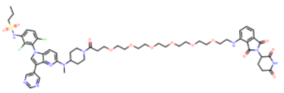 | 83 | 1180 | 3040 | 0.022 | 0.388 | Nat Chem Biol. 2020, 16(11):1170-1178 |
| 1081 | 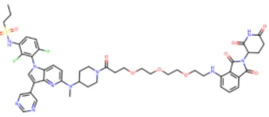 | 65 | 1520 | 3750 | 0.017 | 0.405 | Nat Chem Biol. 2020, 16(11):1170-1178 |
| 339 | 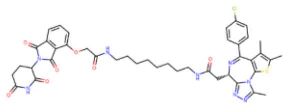 | \ | 800 | 1800/4100 | \ | 0.44 | Nat Chem Biol. 2018, 14(7):706-714 |
| 796 | 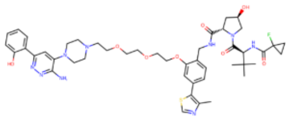 | 2400/600 SMARCA2 | 234/98 | 205/120 | 11.7/5 | 1.14/0.81 | Nat Chem Biol. 2019, 15(7):672-680 |
| 796 | 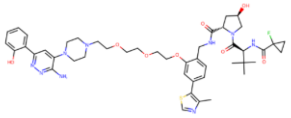 | 3000/1300 SMARCA4 | 234/98 | 230/330 | 13/3.9 | 1.02/0.29 | Nat Chem Biol. 2019, 15(7):672-680 |
| 798 | 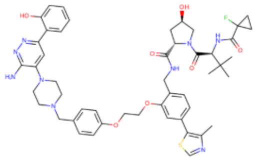 | 770/700 SMARCA2 | 250 | 26/730 | 29.6/0.96 | 9.6/0.34 | Nat Chem Biol. 2019, 15(7):672-680 |
| 798 | 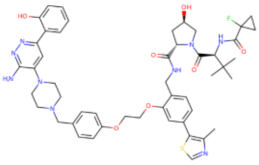 | 560/670 SMARCA4 | 250 | 51/720 | 10.9/0.93 | 4.9/0.35 | Nat Chem Biol. 2019, 15(7):672-680 |
| 797 | 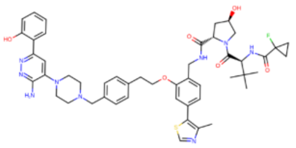 | 790/680 SMARCA2 | 100 | 45/120 | 17.5/5.67 | 2.2/0.83 | Nat Chem Biol. 2019, 15(7):672-680 |

| ID | Structure | $K_a$ (nM) | $K_b$ (nM) | $K_d$ (nM) | $\alpha$ (POI) | $\alpha'$ ($E_3$ ligase) | Quote from |
|---|---|---|---|---|---|---|---|
| 797 | 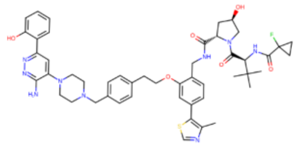 | 990/960 SMARCA4 | 100 | 76/240 | 13.02/4 | 1.31/0.42 | Nat Chem Biol. 2019, 15(7):672-680 |
| 1 | 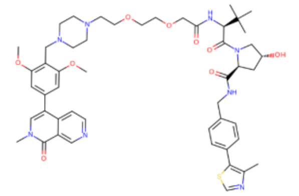 | 15 | 33/24 | 73/98 | 0.2/0.15 | 0.45/0.24 | J Med Chem. 2019, 62(2):699-726 |
| 2389 | 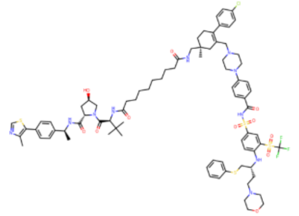 | 2.75 | \ | 0.64 | 4.3 | \ | J Med Chem. 2021, 64(19):14230-14246 |
| 2390 | 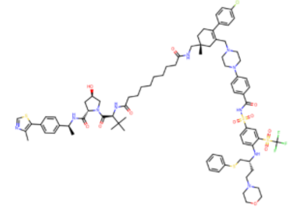 | 2.59 | \ | 0.65 | 3.98 | \ | J Med Chem. 2021, 64(19):14230-14246 |
| 1535 | 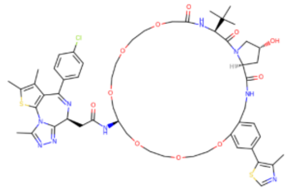 | 180 | 47 | 2 | 90 | 23.5 | Angew Chem Int Ed Engl. 2020,59(4):1727-1734 |
| 2339 | 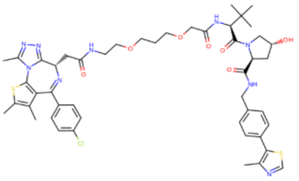 | \ | 66 | 3 | \ | 22 | J Med Chem. 2021, 64(17):12831-12854 |
| 2338 | 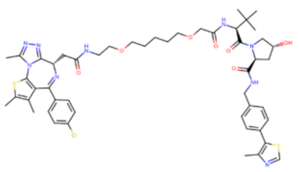 | \ | 98 | 3.7 | \ | 26.5 | J Med Chem. 2021, 64(17):12831-12854 |
| 2630 | 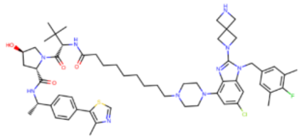 | \ | 56 | 3.7 | \ | 15.1 | J Med Chem. 2022, 655:3923-3942 |

| ID | Structure | $K_a$ (nM) | $K_b$ (nM) | $K_d$ (nM) | $\boldsymbol{\alpha}$ (POI) | $\boldsymbol{\alpha'}$ ($E_3$ ligase) | Quote from |
|---|---|---|---|---|---|---|---|
| 3150 | 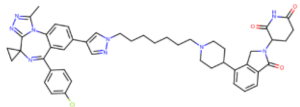 | 58 | 2100 | 4 | 14.5 | 525 | ACS Chem Biol. 2021 16(11):2228-2243 |
| 2340 | 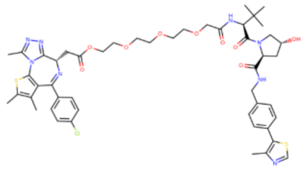 | \ | 248 | 6.4 | \ | 38.75 | J Med Chem. 2021 64(17):12831-12854 |
| 2341 | 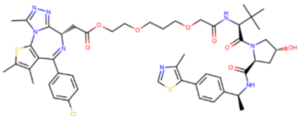 | \ | 63 | 7.3 | \ | 8.63 | J Med Chem. 2021 64(17):12831-12854 |
| 4242 | 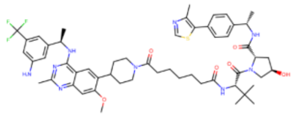 | 93 | \ | 8.3 | 11.2 | \ | J Med Chem. 2023, 666:4197-4214 |
| 335 | 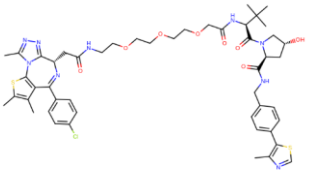 | 26 | 69 | 9 | 2.89 | 7.67 | J Med Chem. 2018, 61(2):504-513 |
| 336 | 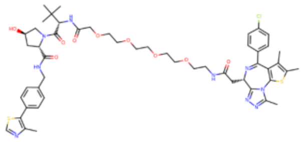 | 27 | 73 | 15 | 1.8 | 4.87 | J Med Chem. 2018, 61(2):504-513 |
| 4251 | 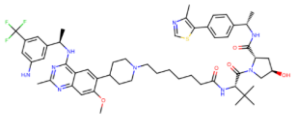 | 44.2 | \ | 16.3 | 2.71 | \ | J Med Chem. 2023, 666:4197-4214 |
| 4247 | 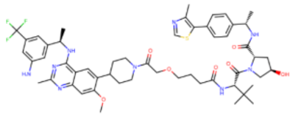 | 155 | \ | 17.4 | 8.91 | \ | J Med Chem. 2023, 666:4197-4214 |
| 2342 | 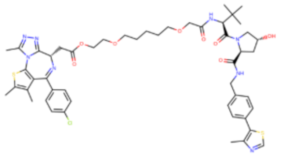 | \ | 236 | 18 | \ | 13.11 | J Med Chem. 2021 64(17):12831-12854 |
| 713 | 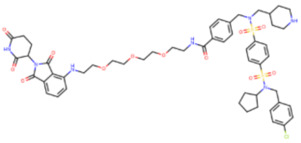 | 27 | \ | 18.4 | 1.47 | \ | Angew Chem Int Ed Engl. 2020, 59(14):5595-5601 |

| ID | Structure | $K_a$ (nM) | $K_b$ (nM) | $K_d$ (nM) | $\alpha$ (POI) | $\alpha'$ ($E_3$ ligase) | Quote from |
|---|---|---|---|---|---|---|---|
| 2343 | 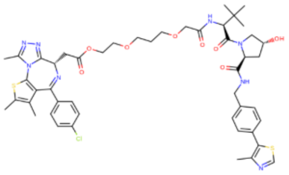 | \ | 143 | 21 | \ | 6.81 | J Med Chem. 2021 64(17):12831-12854 |
| 229 | 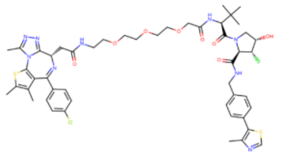 | 28 | 249 | 26 | 1.09 | 9.58 | J Am Chem Soc. 2018, 140(29):9299-9313 |
| 1654 | 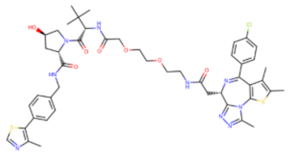 | 17 | 147 | 26 | 0.65 | 5.65 | J Med Chem. 2018, 61(2):504-513 |
| 712 | 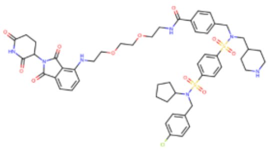 | 14.8 | \ | 26.6 | 0.56 | \ | Angew Chem Int Ed Engl. 2020, 59(14):5595-5601 |
| 4241 | 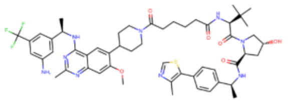 | 75.7 | \ | 26.7 | 2.84 | \ | J Med Chem. 2023, 66(6):4197-4214 |
| 4243 | 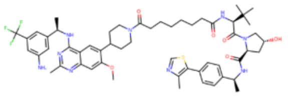 | 78.6 | \ | 27.1 | 2.9 | \ | J Med Chem. 2023, 66(6):4197-4214 |
| 4249 | 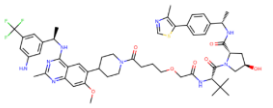 | 63.5 | \ | 27.8 | 2.28 | | J Med Chem. 2023, 66(6):4197-4214 |
| 339 | 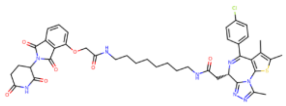 | 46 | 264 | 33 | 1.39 | 8 | ACS Chem Biol. 2021, 16(11):2228-2243 |
| 4248 | 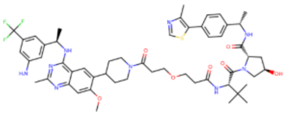 | 95.4 | \ | 33.6 | 2.84 | \ | J Med Chem. 2023, 66(6):4197-4214 |
| 230 | 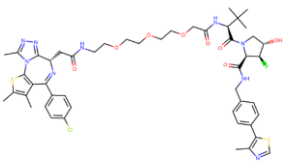 | 32 | 603 | 42 | 0.76 | 14.3 | J Am Chem Soc. 2018, 140(29):9299-9313 |
| 3255 | 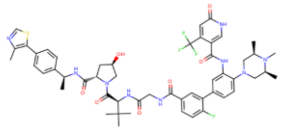 | 63 | 140 | 52 | 1.21 | 2.69 | Sci Transl Med. 2021, 13613:eabj1578 |

| ID | Structure | $K_a$ (nM) | $K_b$ (nM) | $K_d$ (nM) | $\alpha$ (POI) | $\alpha'$ ($E_3$ ligase) | Quote from |
|---|---|---|---|---|---|---|---|
| 5634 | 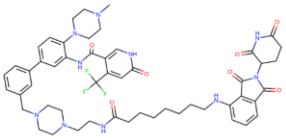 | 125 | 14800 | 63.7 | 1.99 | 232.3 | Oncogene. 2022, 41(24):3328-3340 |
| 229 | 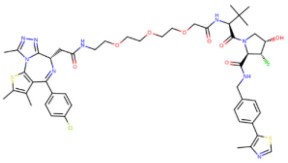 | 97 | 249 | 64 | 1.52 | 3.89 | J Am Chem Soc. 2018, 140(29):9299-9313 |
| 1535 | 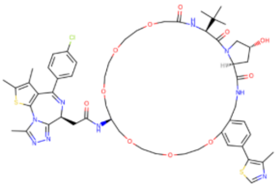 | 743 | 47 | 70 | 10.61 | 0.67 | Angew Chem Int Ed Engl. 2020, 59(4):1727-1734 |
| 714 | 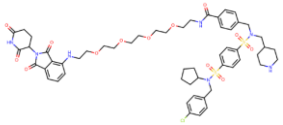 | 64.3 | \ | 92.9 | | \ | Angew Chem Int Ed Engl. 2020, 59(14):5595-5601 |
| 2044 | 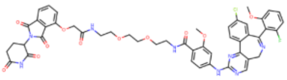 | 375 | 6880 | 183 | 2.05 | 37.59 | Nat Chem Biol. 2020, 16(11):1179-1188 |
| 1653 | 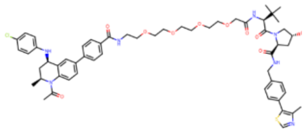 | \ | 69 | 185 | \ | 0.37 | ACS Chem Biol. 2019, 14(3):361-368 |
| 1652 | 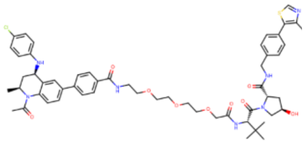 | 4 | 105 | 228 | 0.017 | 0.46 | ACS Chem Biol. 2019, 14(3):361-368 |
| 3254 | 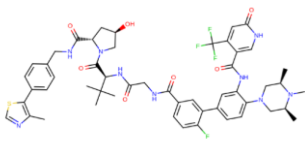 | 122 | 534 | 232 | 0.526 | 2.30 | J Med Chem. 2023 66(23):16168-16186 |
| 2067 | 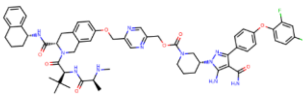 | 692 | \ | 262 | 2.64 | \ | Nat Chem Biol. 2021, 17(2):152-160 |
| 185 | 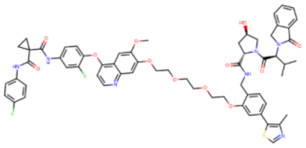 | 550 | 3500 | 436 | 1.26 | 8.03 | Nat Commun. 2019, 10(1):131 |

| ID | Structure | $K_a$ (nM) | $K_b$ (nM) | $K_d$ (nM) | $\alpha$ (POI) | $\alpha'$ ($E_3$ ligase) | Quote from |
|---|---|---|---|---|---|---|---|
| 1651 | 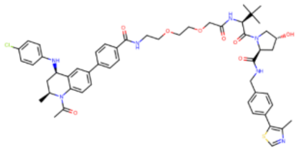 | \ | 104 | 465 | \ | 0.22 | ACS Chem Biol. 2019, 14(3):361-368 |
| 221 | 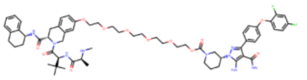 | 259 | 7 | 474 | 0.55 | 0.015 | Nat Chem Biol. 2021, 17(2):152-160 |
| 230 | 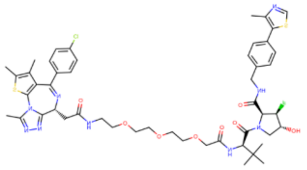 | 111 | 603 | 500 | 0.222 | 1.206 | J Am Chem Soc. 2018, 140(29):9299-9313 |
| 3245 | 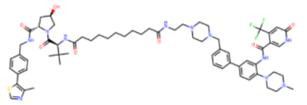 | 120 | 870 | 520 | 0.23 | 1.67 | Sci Transl Med. 2021, 13613:eabj1578 |
| 1651 | 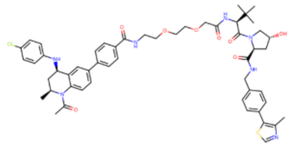 | 3 | 116 | 781 | 0.004 | 0.15 | J Med Chem. 2018, 61(2):504-513 |
| 184 | 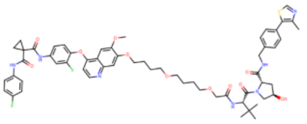 | 500 | 5000 | 1200 | 0.416 | 4.167 | Nat Commun. 2019, 10(1):131 |
| 335 | 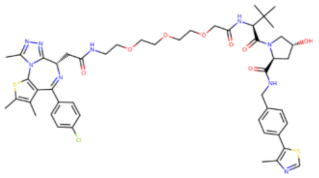 | \ | 29 | 12/8 | \ | 2.42/3.65 | ACS Chem Biol. 2019, 14(3):361-368 |
| 1653 | 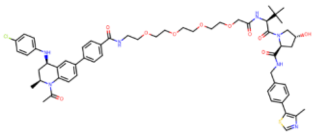 | 39/11 | 109 | 152/434 | 0.256/ 0.025 | 0.717/ 0.251 | J Med Chem. 2018, 61(2):504-513 |
| 335 | 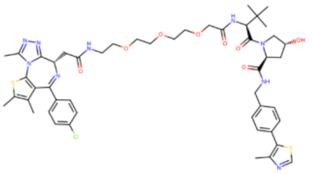 | 21/13 | 66 | 19/7 | 1.1/ 1.85 | 3.47/9.43 | Nat Chem Biol. 2017, 13(5):514-521 |
| 1653 | 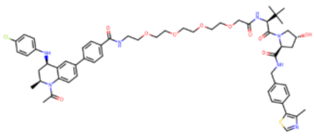 | 39/8 | 109 | 221/183 | 0.17/ 0.04 | 0.49/0.59 | J Med Chem. 2018, 61(2):504-513 |

| ID | Structure | $K_a$ (nM) | $K_b$ (nM) | $K_d$ (nM) | $\boldsymbol{\alpha}$ (POI) | $\boldsymbol{\alpha'}$ ($E_3$ ligase) | Quote from |
|---|---|---|---|---|---|---|---|
| 335 | 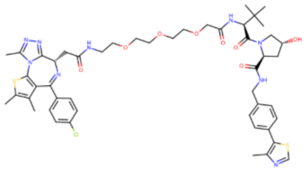 | \ | 29 | 23/0.9 | \ | 1.26/32.2 | Nat Chem Biol. 2017, 13(5):514-521 |
| 335 | 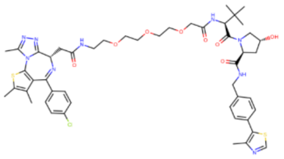 | 62/60 | 66 | 24/28 | 2.58/2.14 | 2.75/2.36 | Nat Chem Biol. 2017, 13(5):514-521 |
| 335 | 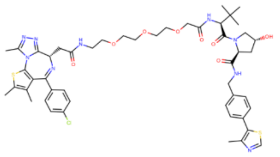 | 39/15 | 66 | 28/3.7 | 1.39/4.05 | 2.36/17.8 | Nat Chem Biol. 2017, 13(5):514-521 |
| 335 | 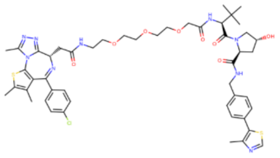 | 62/60 | 29 | 30/1 | 2.06/60 | 0.97/29 | Nat Chem Biol. 2017, 13(5):514-521 |
| 22 | 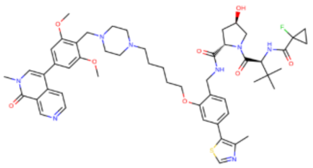 | 5.1 | 26/35 | 27/35 | 0.189/0.15 | 0.96/1 | J Med Chem. 2019, 62(2):699-726 |
| 1653 | 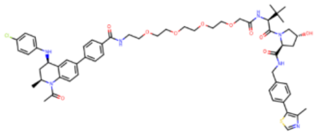 | 45/5 | 109 | 328/384 | 0.14/0.013 | 0.33/0.28 | J Med Chem. 2018, 61(2):504-513 |
| 310 | 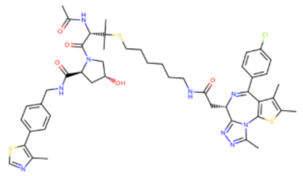 | 75/44 | 335 | 390/46 | 0.19/0.96 | 0.86/7.28 | Nat Chem Biol. 2017, 13(5):514-521 |
| 310 | 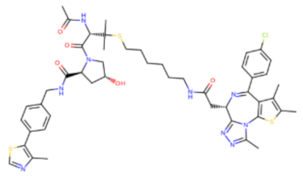 | \ | 110 | 578/24 | \ | 0.19/4.58 | Nat Chem Biol. 2017, 13(5):514-521 |
| 11 | 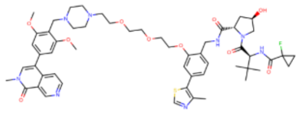 | 8.8 | 87/70 | 83/60 | 0.11/0.15 | 1.05/1.17 | J Med Chem. 2019, 62(2):699-726 |

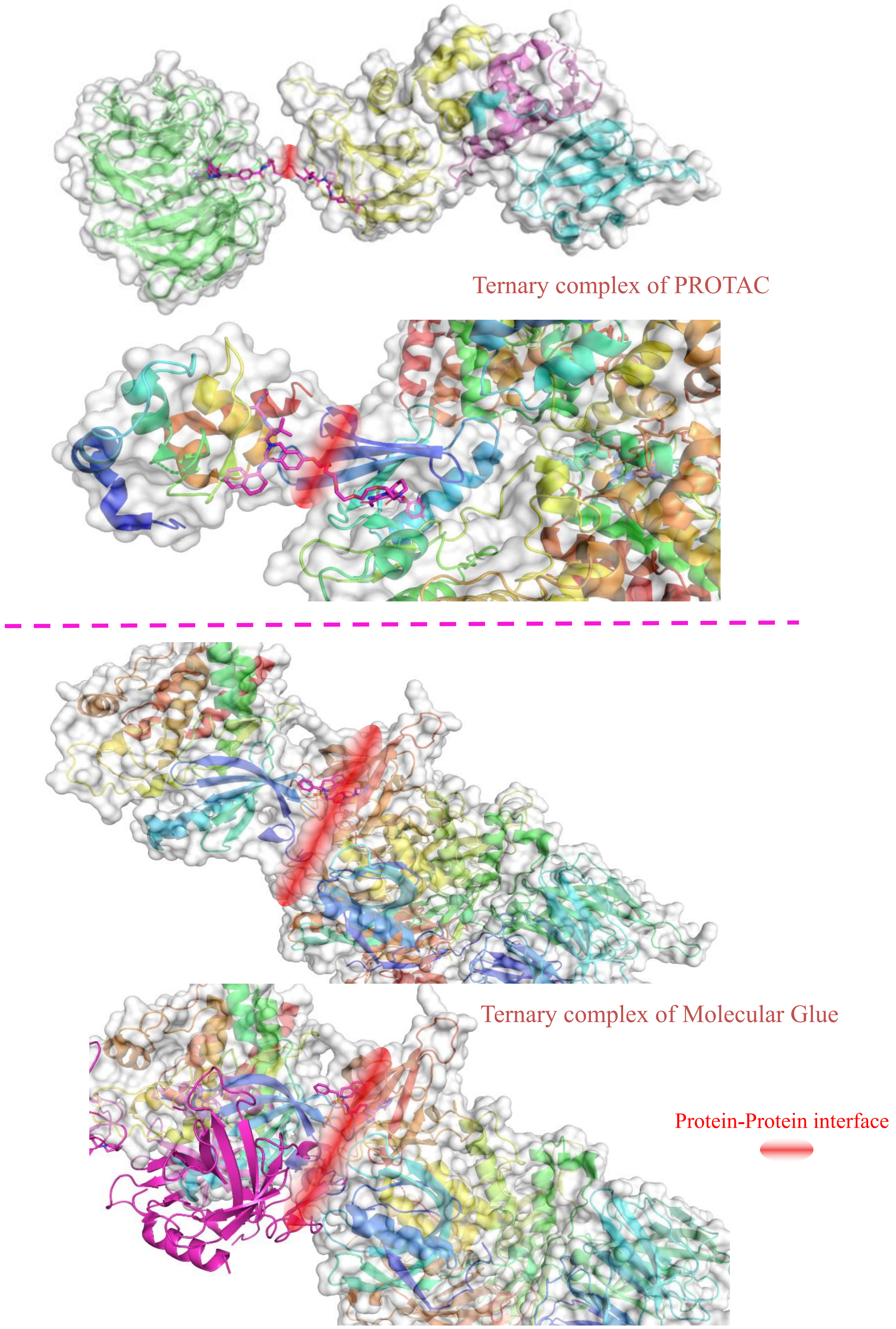


**Figure S1**. Cryo-EM of Ternary complex structure of PROTAC (PDB:8BB2 and 6W8I) and Ternary complex structure of Molecular Glue (PDB:9NFQ and 909I).

Figure S2. Cases of the report $IC_{50}$ value for the warhead and its related PROTAC molecules.

$IC_{50}$(CDK2) = 0.7 nM

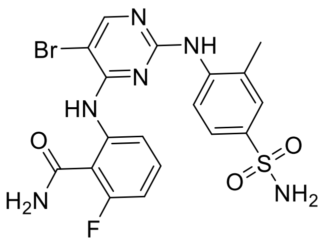

PROTAC(TMX-2172)
$IC_{50}$(CDK2) = 6.5 nM

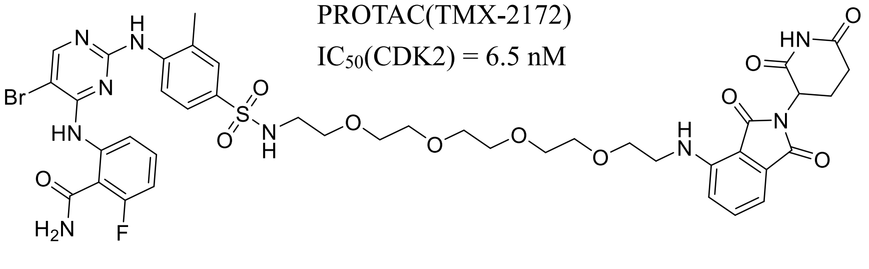

$IC_{50}$(c-Kit) = 0.064 nM

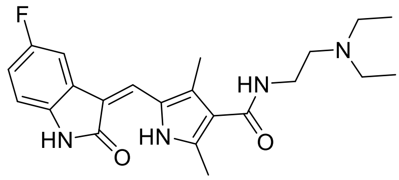

PROTAC(MI-389)
$IC_{50}$(c-Kit) = 0.55 nM

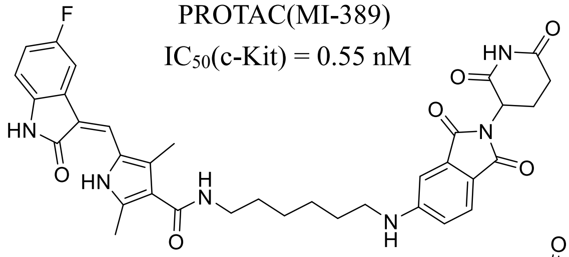

$IC_{50}$(CDK4) = 10 nM

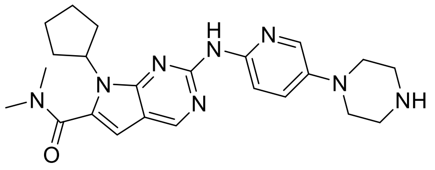

PROTAC(BSJ-04-132)
$IC_{50}$(CDK4) = 50.6 nM

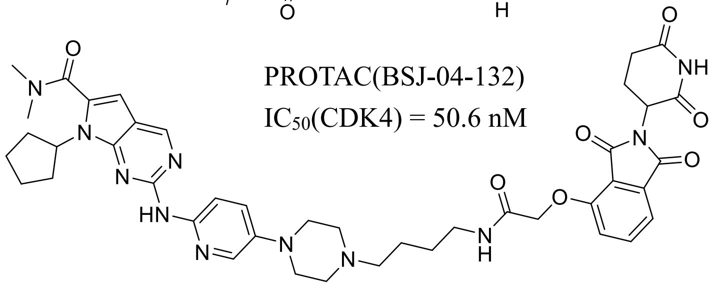

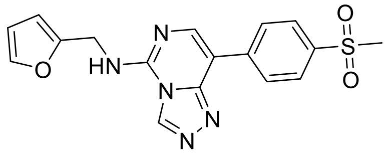

$IC_{50}$(EED) = 22 nM

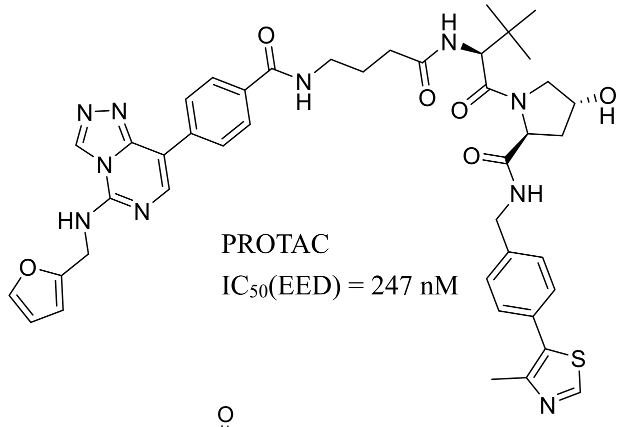

PROTAC
$IC_{50}$(EED) = 247 nM

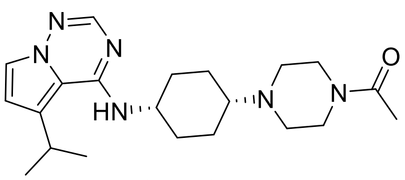

$IC_{50}$(IRK3) = 6 nM

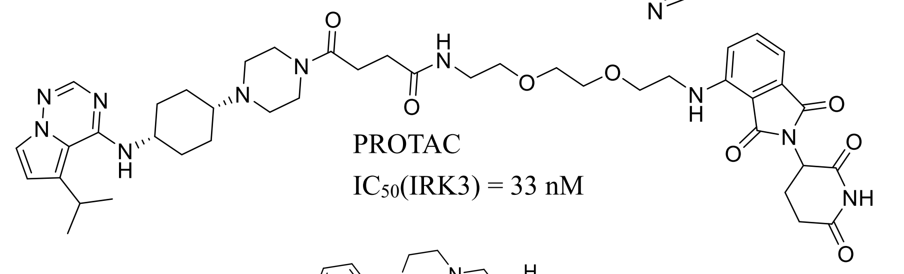

PROTAC
$IC_{50}$(IRK3) = 33 nM

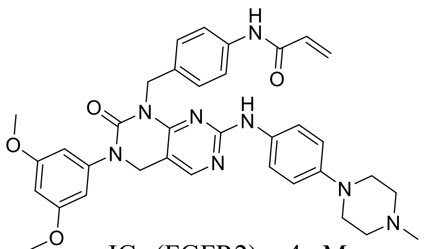

$IC_{50}$(FGFR2) = 4 nM

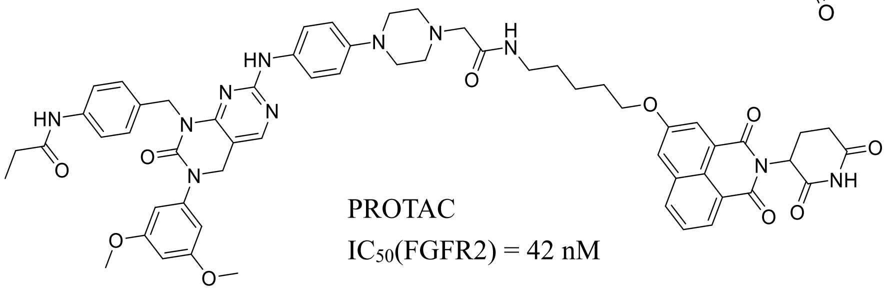

PROTAC
$IC_{50}$(FGFR2) = 42 nM

Figure S3. Cases of the report $IC_{50}$ value of the PROTAC molecules with different length unit's linker.

SARS-CoV-2 Degrader

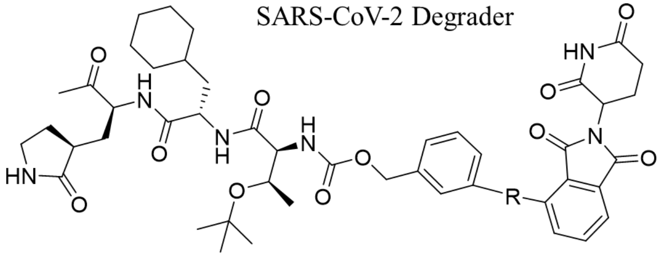

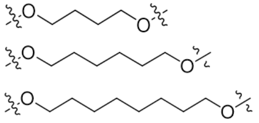

| | | | |
|---|---|---|---|
| MPD1 | R= | M.W. 959.48 | IC50 = 39 nM |
| MPD2 | R= | M.W. 983.53 | IC50 = 41 nM |
| MPD3 | R= | M.W. 1007.58 | IC50 = 51 nM |

BRD9 Degrader

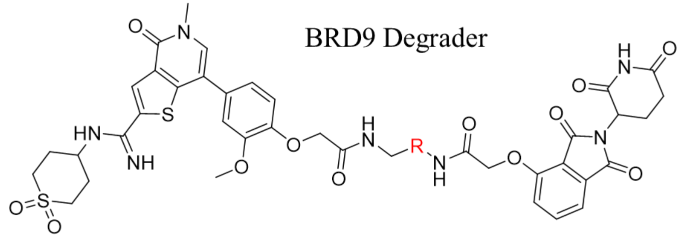

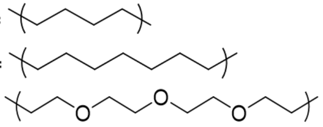

| | | | |
|---|---|---|---|
| 1187 | R= | M.W. 904.0 | $IC_{50}$ = 13.5 nM |
| 1188 | R= | M.W. 960.09 | $IC_{50}$ = 18.9 nM |
| 1189 | R= | M.W. 1036.14 | $IC_{50}$ = 36.5 nM |

BRD9 Degrader

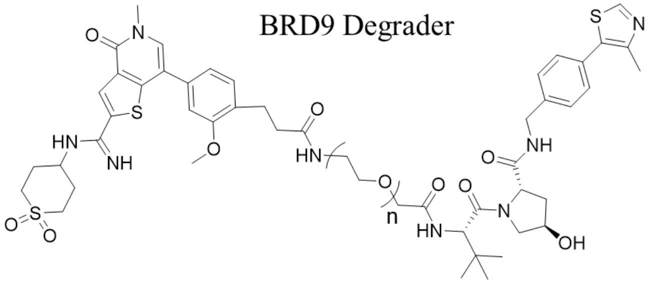

| | | | |
|---|---|---|---|
| 1193 | n=2 | M.W. 1073.37 | $IC_{50}$ = 22.7 nM |
| 1199 | n=3 | M.W. 1117.42 | $IC_{50}$ = 43.2 nM |

CDK2 Degrader

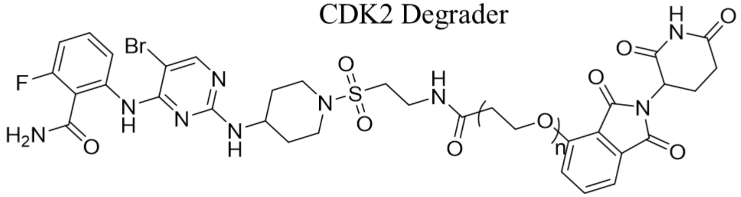

| | | | |
|---|---|---|---|
| TMX-1128 | n=3 | M.W. 932.79 | $IC_{50}$ = 4.9 nM |
| TMX-1111 | n=5 | M.W. 1020.91 | $IC_{50}$ = 6.4 nM |
| TMX-1146 | n=7 | M.W. 1109.03 | $IC_{50}$ = 7.3 nM |

PARP2 Degrader

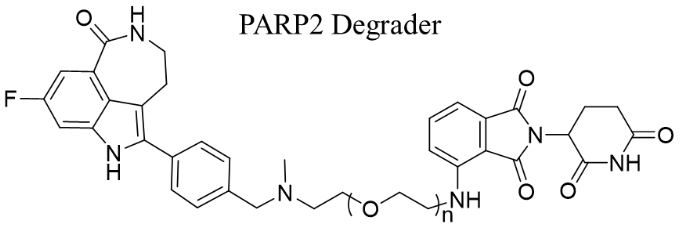

| | | | |
|---|---|---|---|
| iRucaparib-AP5 | n=5 | M.W. 941.59 | $IC_{50} = 5.1 \pm 0.2$ nM |
| iRucaparib-AP6 | n=6 | M.W. 985.59 | $IC_{50} = 7.2 \pm 0.3$ nM |

CDK2 Degrader

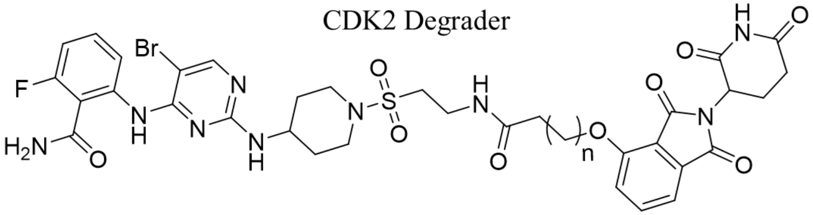

| | | | |
|---|---|---|---|
| TMX-1162 | n=6 | M.W. 914.81 | $IC_{50}$ = 17.1 nM |
| TMX-1160 | n=7 | M.W. 983.53 | $IC_{50}$ = 28.7 nM |

#### PARP1 Degrader

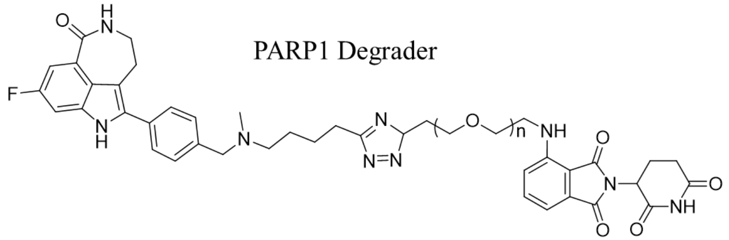

| | | | |
|---|---|---|---|
| iRucaparib-TP1 | n=1 | M.W. 888.87 | $IC_{50}$ = 13.35 nM |
| iRucaparib-TP2 | n=2 | M.W. 932.94 | $IC_{50}$ = 14.00 nM |
| iRucaparib-TP3 | n=3 | M.W. 977.01 | $IC_{50}$ = 18.26 nM |
| iRucaparib-TP4 | n=4 | M.W. 1021.08 | $IC_{50}$ = 31.81 nM |

#### APT1 Degrader

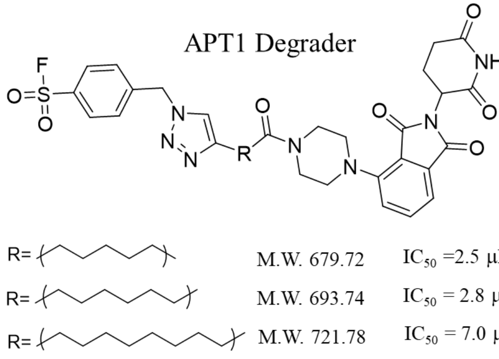

| | | |
|---|---|---|
| R= (linker 1) | M.W. 679.72 | $IC_{50}$ =2.5 μM |
| R= (linker 2) | M.W. 693.74 | $IC_{50}$ = 2.8 μM |
| R= (linker 3) | M.W. 721.78 | $IC_{50}$ = 7.0 μM |

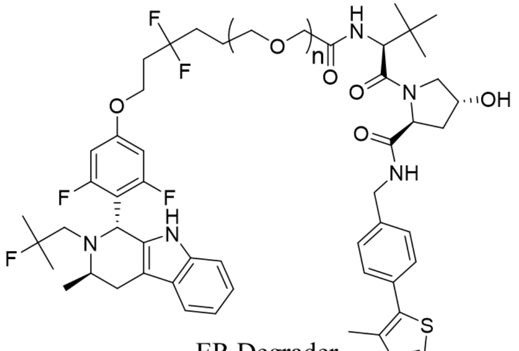

#### ER Degrader

| | | | |
|---|---|---|---|
| 1469 | n=2 | M.W. 1025.19 | $IC_{50}$ = 0.5 nM |
| 1470 | n=3 | M.W. 1069.24 | $IC_{50}$ = 0.6 nM |
| 1471 | n=4 | M.W. 1113.29 | $IC_{50}$ = 0.7 nM |

#### BRAF V600E Degrader

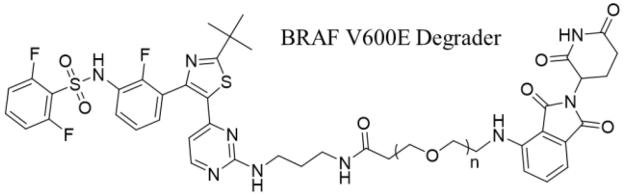

| | | |
|---|---|---|
| n=3 | M.W. 1036.11 | $IC_{50}$ = 99 nM |
| n=4 | M.W. 1080.18 | $IC_{50}$ = 122 nM |
| n=6 | M.W. 1168.32 | $IC_{50}$ = 199 nM |

#### BRD4 Degrader

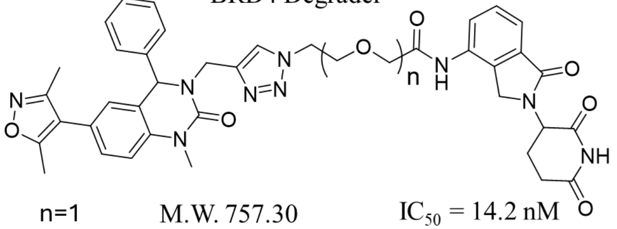

| | | |
|---|---|---|
| n=1 | M.W. 757.30 | $IC_{50}$ = 14.2 nM |
| n=2 | M.W. 801.37 | $IC_{50}$ = 15.4 nM |
| n=3 | M.W. 845.44 | $IC_{50}$ = 22.1 nM |

#### EZH2 Degrader

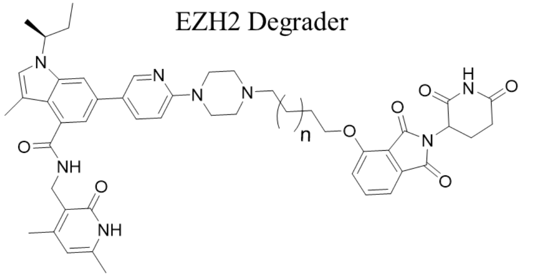

| | | |
|---|---|---|
| n=1 | M.W. 855.01 | $IC_{50}$ = 5.5 nM |
| n=2 | M.W. 869.04 | $IC_{50}$ = 5.8 nM |
| n=4 | M.W. 897.11 | $IC_{50}$ = 6.3 nM |
| n=5 | M.W. 911.13 | $IC_{50}$ = 7.3 nM |

#### CYP1B1 Degrader

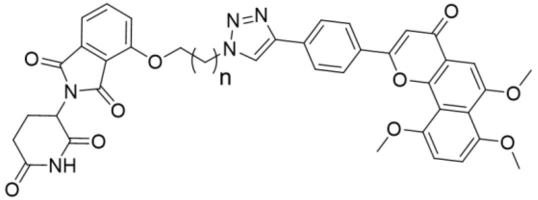

| | | |
|---|---|---|
| n=1 | M.W. 729.21 | $IC_{50}$ = 101.2 nM |
| n=4 | M.W. 771.80 | $IC_{50}$ = 152.7 nM |
| n=7 | M.W. 813.88 | $IC_{50}$ = 166.2 nM |

#### FAK Degrader

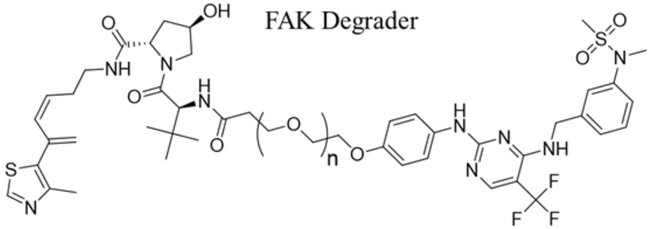

| | | |
|---|---|---|
| n=1 | M.W. 986.14 | $IC_{50}$ = 6.5 nM |
| n=2 | M.W. 1030.21 | $IC_{50}$ = 12.7 nM |

**Table S4**. The prediction value of $IC_{50}$ or $K'_b$ using the Ligand entropy method simulation compared with commercial software.

| Report data | | | | Pred. by ligand entropy method | | | Pred. by MOE 2019 | | | Pred. by Schordinger Maestro 13.3 | | |
|---|---|---|---|---|---|---|---|---|---|---|---|---|
| Comp. NO. | $IC_{50}$ (nM) | $K'b$ (μM) | ΔG (KJ/mol) | $IC_{50}$ (nM) | $K'b$ (μM) | ΔG (KJ/mol) | $IC_{50}$ (nM) | $K'b$ (μM) | ΔG (KJ/mol) | $IC_{50}$ (nM) | $K'b$ (μM) | ΔG (KJ/mol) |
| Warhead [1] (CRBN) | 140 | | -39.11 | (140) | | – | 34480 | | -25.46 | 14000 | | -27.69 |
| 16 | | 0.9±0.04 | -34.49 | | 0.898 | -34.49 | | 0.112 | -39.66 | | 0.923 | -34.44 |
| 32 | | 1.3±0.05 | -33.58 | | 1.28 | -33.58 | | 0.112 | -39.66 | | 0.057 | -41.33 |
| 33 | | 1.8±0.00 | -32.77 | | 1.19 | -33.78 | | 0.6 | -35.51 | | 0.247 | -37.70 |
| 34 | | 2.8±0.14 | -31.68 | | 1.19 | -33.78 | | 0.035 | -42.57 | | 0.021 | -43.81 |
| 35 | | 1.3±0.01 | -33.58 | | 1.28 | -33.58 | | 0.74 | -34.98 | | 0.872 | -34.57 |
| 36 | | 1.2±0.01 | -33.78 | | 1.19 | -33.78 | | 1.55 | -33.15 | | 0.447 | -36.23 |
| 37 | | 1.1±0.03 | -33.99 | | 1.11 | -33.97 | | 0.345 | -36.87 | | 0.05 | -41.65 |
| Warhead [2] (Rucaparib) | 2.46 | | -49.12 | (2.46) | | – | 3670 | | -31.01 | 5023 | | -30.23 |
| iRucaparib-TP1 | 13.4±2.1 | | -44.92 | 13.72 | | -44.86 | 4.178 | | -47.81 | 32.36 | | -42.74 |
| iRucaparib-TP2 | 14.0±1.6 | | -44.81 | 15.04 | | -44.64 | 0.3303 | | -54.09 | 69.5 | | -40.84 |
| iRucaparib-TP3 | 18.3±2.2 | | -44.15 | 16.41 | | -44.42 | 0.0178 | | -61.34 | 42.85 | | -42.04 |
| iRucaparib-TP4 | 31.8±3.2 | | -42.78 | 30.11 | | -42.91 | 0.1175 | | -56.66 | 16.75 | | -44.37 |
| iRucaparib-AP5 | 24.6±7.2 | | -43.42 | 25.15 | | -43.36 | 0.1079 | | -56.87 | 2.63 | | -48.96 |
| iRucaparib-AP6 | 28.5±7.4 | | -43.05 | 27.85 | | -43.10 | 0.1413 | | -56.20 | 0.85 | | -51.75 |

Table S5. The original ITC thermodynamics cycle data of binary/ternary complex from literature.[3]

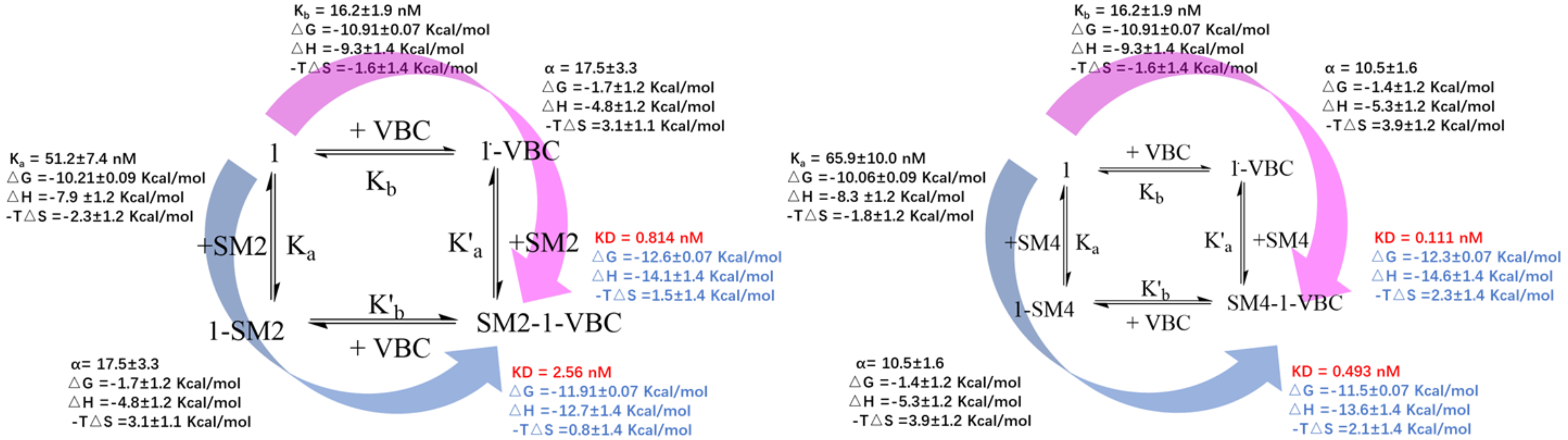


| PROTAC ($M$) | POI ($M$) | $K_a$ $\Delta H$ (Kcal/mol) | $K_a - T\Delta S$ (Kcal/mol) | $K'_a$ $\Delta H'$ (Kcal/mol) | $K'_a - T\Delta S'$ (Kcal/mol) | $\Delta\Delta H$ (Kcal/mol) | $-T\Delta\Delta S$ (Kcal/mol) | $\Delta\Delta G$ (Kcal/mol) | $\Delta S_{ligand}$ (Kcal/mol) | $\log[\alpha]_{PPI}$ (Kcal/mol) | $\Delta S_{Conf}$ (Kcal/mol) | $\Delta S_{Comp}$ (Kcal/mol) |
|---|---|---|---|---|---|---|---|---|---|---|---|---|
| Comp.1 (819) | SM2 (14.3KDa) | -7.9±1.2 | -2.3±1.2 | -4.8±1.2 | 3.1±1.1 | +3.1 | +5.4 | 8.5 | +5.972 | +2.528 | -0.572 | +3.1 |
| Comp.1 (819) | SM4 (14.7KDa) | -8.3±1.2 | -1.8±1.2 | -5.3±1.2 | 3.9±1.2 | +3.0 | +5.7 | 8.7 | +5.972 | +2.738 | -0.272 | +3.0 |

| Titration path | $\Delta G_{Total}$ (Kcal/mol) | $\Delta H_{Total}$ (Kcal/mol) | $-T\Delta S_{Total}$ (Kcal/mol) | Titration path | $\Delta G_{Total}$ (Kcal/mol) | $\Delta H_{Total}$ (Kcal/mol) | $-T\Delta S_{Total}$ (Kcal/mol) |
|---|---|---|---|---|---|---|---|
| 1 + VBC→1• VBC<br>1• VBC + SM2 → SM2•1• VBC | -12.6 | -14.1 | 1.5 | 1 + SM2 →1• SM2<br>1• SM2 + VBC → SM2•1• VBC | -11.9 | -12.7 | 0.8 |
| 1 + VBC→1• VBC<br>1• VBC + SM4 → SM4•1• VBC | -12.3 | -14.6 | 2.3 | 1 + SM4 →1• SM4<br>1• SM4 + VBC → SM4•1• VBC | -11.5 | -13.6 | 2.1 |

Table S6. Revised full ITC thermodynamic data of binary/ternary complex from literature.[3]

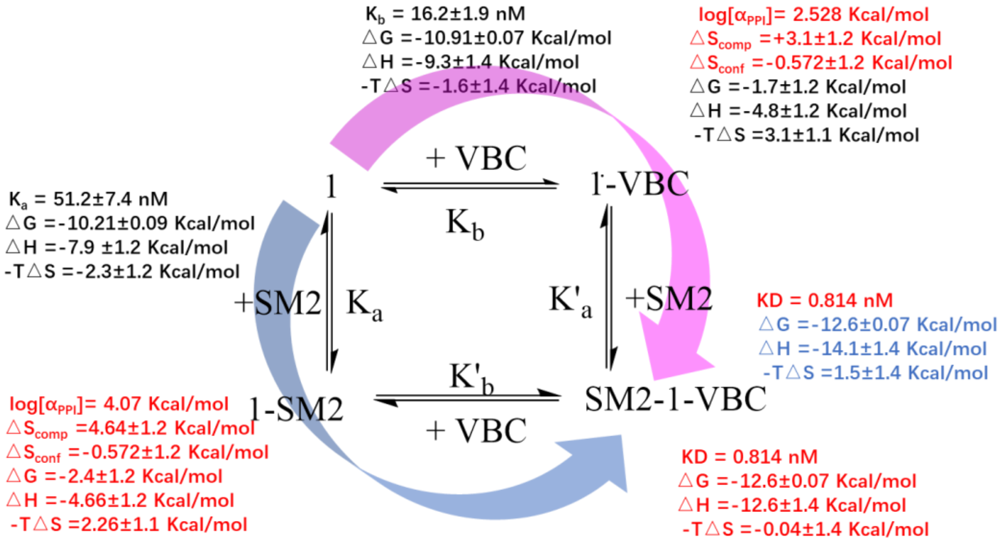


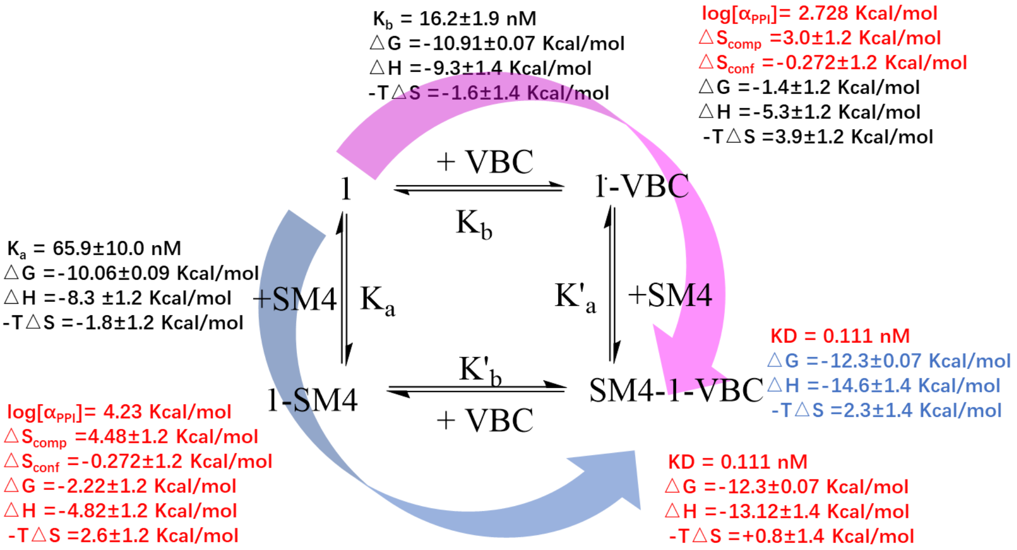


| | | | | | | | | | | | |
|---|---|---|---|---|---|---|---|---|---|---|---|
| Clockwise SM2 | $K_a$ $\Delta H$ (Kcal/mol) | $K_a - T\Delta S$ (Kcal/mol) | $K'_a$ $\Delta H'$ (Kcal/mol) | $K'_a - T\Delta S'$ (Kcal/mol) | $\Delta\Delta H$ (Kcal/mol) | $-T\Delta\Delta S$ (Kcal/mol) | $\Delta\Delta G$ (Kcal/mol) | $\Delta S_{ligand}$ (Kcal/mol) | $\log[\alpha]_{PPI}$ (Kcal/mol) | $\Delta S_{Conf}$ (Kcal/mol) | $\Delta S_{Comp}$ (Kcal/mol) |
| | -7.9±1.2 | -2.3±1.2 | -4.8±1.2 | 3.1±1.1 | +3.1 | +5.4 | +8.5 | +5.972 | +2.528 | -0.572 | +3.1 |
| Anti-Clock SM2 | $K_b$ $\Delta H$ | $K_b - T\Delta S$ | $K'_b$ $\Delta H'$ | $K'_b - T\Delta S'$ | $\Delta\Delta H$ | $-T\Delta\Delta S$ | $\Delta\Delta G$ | $\Delta S_{ligand}$ | $\log[\alpha]_{PPI}$ | $\Delta S_{Conf}$ | $\Delta S_{Comp}$ |
| | -9.3±1.2 | -1.6±1.2 | +4.66±1.2 | 2.26±1.2 | +4.64 | +3.86 | +8.5 | +4.43 | +4.07 | -0.572 | +4.64 |
| Clockwise SM4 | $K_a$ $\Delta H$ | $K_a - T\Delta S$ | $K'_a$ $\Delta H'$ | $K'_a - T\Delta S'$ | $\Delta\Delta H$ | $-T\Delta\Delta S$ | $\Delta\Delta G$ | $\Delta S_{ligand}$ | $\log[\alpha]_{PPI}$ | $\Delta S_{Conf}$ | $\Delta S_{Comp}$ |
| | -8.3±1.2 | -1.8±1.2 | -5.3±1.2 | 3.9±1.2 | +3.0 | +5.7 | +8.7 | +5.972 | +2.728 | -0.272 | +3.0 |
| Anti-Clock SM4 | $K_b$ $\Delta H$ | $K_b - T\Delta S$ | $K'_b$ $\Delta H'$ | $K'_b - T\Delta S'$ | $\Delta\Delta H$ | $-T\Delta\Delta S$ | $\Delta\Delta G$ | $\Delta S_{ligand}$ | $\log[\alpha]_{PPI}$ | $\Delta S_{Conf}$ | $\Delta S_{Comp}$ |
| | -9.3±1.2 | -1.6±1.2 | -4.8±1.2 | 2.6±1.2 | +4.5 | +4.20 | +8.7 | +4.47 | +4.23 | -0.272 | +4.5 |

### REFERENCE


1 Chen, Z. *et al.* Discovery of CBPD-268 as an Exceptionally Potent and Orally Efficacious CBP/p300 PROTAC Degrader Capable of Achieving Tumor Regression. *J Med Chem* **67**, 5275-5304, doi:10.1021/acs.jmedchem.3c02124 (2024).

2 Wang, S. *et al.* Uncoupling of PARP1 trapping and inhibition using selective PARP1 degradation. *Nat Chem Biol* **15**, 1223-+, doi:10.1038/s41589-019-0379-2 (2019).

3 Wurz, R. P. *et al.* Affinity and cooperativity modulate ternary complex formation to drive targeted protein degradation. *Nat Commun* **14**, 4177, doi:10.1038/s41467-023-39904-5 (2023).
